# Changes in Environmental Stress over COVID-19 Pandemic Likely Contributed to Failure to Replicate Adiposity Phenotype Associated with *Krtcap3*

**DOI:** 10.1101/2023.03.15.532439

**Authors:** Alexandria M Szalanczy, Gina Giorgio, Emily Goff, Osborne Seshie, Michael Grzybowski, Jason Klotz, Aron M Geurts, Eva E Redei, Leah C Solberg Woods

**Author notes:** Correspondence: Leah C Solberg Woods. **Funding**. R01 DK106386, T32 DA041349, R01 DK120667.

## Abstract

We previously identified *Keratinocyte-associated protein 3, Krtcap3,* as an obesity-related gene in female rats where a whole-body *Krtcap3* knock-out (KO) led to increased adiposity compared to wild-type (WT) controls when fed a high-fat diet (HFD). We sought to replicate this work to better understand the function of *Krtcap3* but were unable to reproduce the adiposity phenotype. In the current work, WT female rats ate more compared to WT in the prior study, with corresponding increases in body weight and fat mass, while there were no changes in these measures in KO females between the studies. The prior study was conducted before the COVID-19 pandemic, while the current study started after initial lock-down orders and was completed during the pandemic with a generally less stressful environment. We hypothesize that the environmental changes impacted stress levels and may explain the failure to replicate our results. Analysis of corticosterone (CORT) at euthanasia showed a significant study by genotype interaction where WT had significantly higher CORT relative to KO in Study 1, with no differences in Study 2. These data suggest that decreasing *Krtcap3* expression may alter the environmental stress response to influence adiposity. We also found that KO rats in both studies, but not WT, experienced a dramatic increase in CORT after their cage mate was removed, suggesting a separate connection to social behavioral stress. Future work is necessary to confirm and elucidate the finer mechanisms of these relationships, but these data indicate the possibility of *Krtcap3* as a novel stress gene.

## Introduction

Obesity rates continue to climb in the United States. The 2017-2020 pre-pandemic National Health and Nutrition Examination Survey found that rates of childhood obesity were just under 20%, and adult obesity just over 40% (1). But early studies during the COVID-19 pandemic examining obesity rates in children and adults showed that obesity prevalence increased about 3% in each group in just one year alone (2, 3). The obesity epidemic continues to accelerate, and this highlights the severe need in the medical community to better understand the underlying causes of obesity and develop more effective treatment options for patients.

Obesity is a complex disease, influenced by both genetics and environment. Although environmental components such as physical activity, socioeconomic status, and stress exposure are well-recognized (4), the genetic susceptibility to obesity remains poorly understood (5–7). Hundreds of genetic loci for obesity have been identified in human genome wide association studies (GWAS), but they still only explain a small portion of the heritability (6–8). Furthermore, many of the causal genes within these loci are unknown. Notably, most common obesity variants will impact weight subtly, so as more variants are found researchers should expect diminishing effect sizes (9). While there is much work to be done in understanding the genetics of obesity and no gene alone may be a cure-all for patients, it is important to continue to advance the field to improve treatment options and personalize approaches for patients suffering with obesity and its related complications.

Previously, we identified *Keratinocyte-associated protein 3* (*Krtcap3*) as a candidate gene for visceral fat mass in rats (10, 11). At the same time, a different group found that *Krtcap3* may be a pleiotropic gene for obesity, type 2 diabetes, and dyslipidemia in humans (12). Our lab conducted the first published study to investigate *Krtcap3* as an adiposity gene and were able to validate these GWAS findings (13). Specifically, we developed a *Krtcap3* knock-out (KO) rat model on the inbred Wistar-Kyoto background strain. We showed that KO rats had increased body weight in both sexes relative to wild-type (WT) rats, and that female KO rats had increased eating and fat mass relative to WT rats (13). Despite this validation, very little is known about the underlying molecular mechanisms of how *Krtcap3* impacts adiposity.

The initial purpose of the current work was to conduct a second study in order to replicate our previous findings and to probe the functional aspect of *Krtcap3* in obesity. We chose to focus on female rats in this study in order to better understand the role of *Krtcap3,* and then later apply that knowledge to studies with male rats. Surprisingly, we failed to replicate the adiposity difference we had previously seen in the female rats, as there were no longer any significant differences in body weight, food intake, or fat mass between WT and KO females. Vitally, between the first study (Study 1) and this second study (Study 2) there was a large environmental change due to the COVID-19 shutdown and the completion of a nearby construction project. Our results support the importance of environmental factors in contributing to gene by environment interactions and suggest that *Krtcap3* may interact in the stress response pathway, thereby indirectly affecting adiposity in rats.

## Methods

### Animals

As previously described, we generated a whole-body *in vivo Krtcap3* knock-out (KO) on the Wistar-Kyoto (WKY/NCrl; RGD_1358112) inbred rat strain (WKY-Krtcap3^em3Mcwi^) and established a breeding colony at Wake Forest University School of Medicine (WFSoM) in 2019 (13). Rats were housed in the same conditions previously described (13). Rats were housed in standard caging at 22°C in a 12 h light and 12 h dark cycle (dark from 18:00 to 6:00) at standard temperature and humidity conditions, and given *ad libitum* access to water. The breeding colony has been maintained at WFSoM since 2019, with WT and KO experimental rats weaned from heterozygous breeders. Experimental rats for Study 1 (13) were weaned from July 2019 through February 2020, during which time the building was fully occupied and there was a neighboring construction project. Experimental rats for Study 2 (described below) were weaned from May 2020 through November 2020, when the facility operated at minimum capacity and construction had finished. Breeders and experimental rats prior to study start were given standard chow diet (Lab Diet, Prolab RMH 3000, Catalog #5P00). At study start, experimental WT (n = 11) and KO (n = 18) rats were placed on experimental diet as described below.

To conduct analyses of *Krtcap3* expression in multiple tissues, we ordered WKY/NCrl rats from Charles River (male n = 10; female n = 3). Rats remained at WFSoM for at least two weeks in the same housing conditions as described above, on standard chow diet and housed two to three rats per cage. At 12 weeks of age the rats were euthanized and their tissues dissected for expression analyses. Tissues were collected from multiple groups of males from 2019-2022 while tissues were collected from only one group of females in 2022.

### Genotyping

Experimental rats were genotyped by fluorescent based fragment analysis at MCW using the ABI 3730 capillary sequencer, followed by analysis in Genemapper software.

### Study Design

Female WT and KO experimental rats were weaned at three weeks of age and placed two per cage in same-sex, same-genotype cages. We housed WT rats separately from KO rats unless a same genotype cage-mate was not available. Prior to starting a high fat diet (HFD), as described below, experimental rats were maintained on the same diet as breeders (see above). At six weeks of age, rats were weighed, analyzed by EchoMRI analysis, and began a HFD (60% kcal fat; ResearchDiet D12492). Rats were allowed access to diet *ad libitum* with body weight and cage food intake recorded weekly starting at six weeks of age. Weekly caloric intake and cumulative caloric intake were calculated as previously described (13). Rats were on diet for 13 weeks, with metabolic phenotyping tests beginning after 10 weeks on diet and euthanasia after 13 weeks on diet (**Figure 1**). All experiments were performed using a protocol approved by the Institutional Animal care and Use Committee at WFSoM.

**Figure 1.**
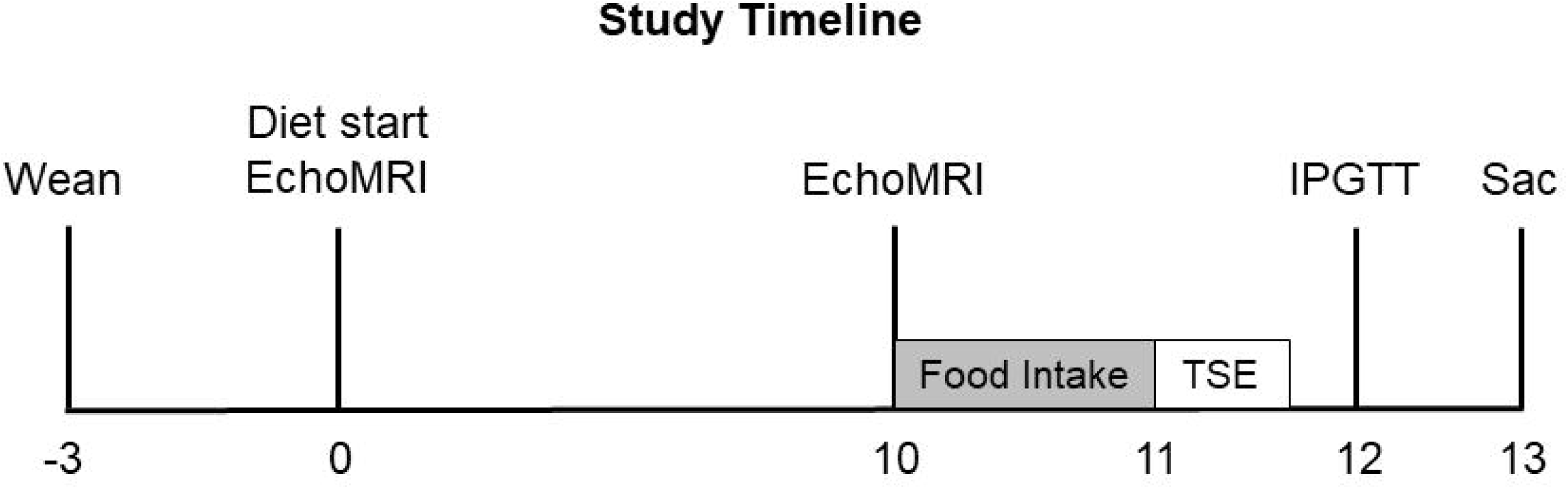
Study timeline. Timeline outlining study design, with weeks relative to diet start shown. Metabolic phenotyping included EchoMRI analysis (EchoMRI), individual food intake (Food Intake, shaded gray), TSE metabolic phenotyping systems (TSE), an intraperitoneal glucose tolerance test (IPGTT), and euthanasia (Sac). Body weight and cage-wide food intake were recorded weekly.

### Metabolic phenotyping

At 0 and 10 weeks on diet, rats went through EchoMRI (EchoMRI LLC, Houston, TX) analysis as previously described (13) to precisely measure total fat and lean mass of live rats.

Immediately following the second EchoMRI analysis, rats began a five-to-seven day-long period of individual housing to measure individual food intake and to measure metabolism. To measure individual food intake, rats were housed in the same standard caging, with the addition of a tube for enrichment, and still allowed access to diet *ad libitum.* Body weight was recorded at the start and the end, and food remaining was recorded every day between 11:00-13:00.

Rats were then transferred to TSE Phenomaster chambers (TSE Systems, Berlin, Germany) to measure metabolism. TSE Phenomaster chambers use indirect gas calorimetry to measure volume of oxygen consumed by the rat and volume of carbon dioxide produced. The system’s climate chamber controls temperature (22 °C), humidity, and lighting conditions, which are held constant to the rat’s home environment. Rats were given a 24 h adaptation period, and then data were recorded for at least an additional 24 h. Total energy expenditure (TEE) was calculated by the indirect calorimetry chamber and normalized by individual body weight. Respiratory exchange ratio (RER), a measure of fuel utilization, was calculated by dividing volume carbon dioxide produced (VCO2) by volume oxygen used (VO_2_). Food intake was also measured during this time period.

After 12 weeks on diet, rats were fasted 16 h overnight before being administered an intraperitoneal glucose tolerance test (IPGTT). Rats were transferred to a procedure room away from the housing room 30 minutes before the test began. We measured blood glucose (Contour Next EZ) and collected serum (Sarstedt Inc MicrovetteCB 300 LH) for subsequent insulin analysis from a tail nick at fasting and 15, 30, 60, 90, and 120 minutes after a 1 mg/kg glucose injection. To calculate a rat’s response to the glucose challenge, the glucose area-under-the-curve (AUC), we used the following equation:

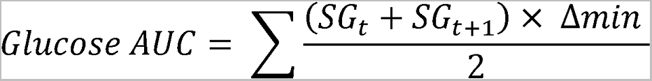

Where serum glucose (SG) was measured at t = 0, 15, 30, 60, 90, and 120 min.

### Tissue harvest

After 13 weeks on diet, rats were euthanized via decapitation after either no fast or a 4 h morning fast. Statistical analysis confirmed that length of fast did not affect adiposity measures. Rats were transferred to the anteroom of the necropsy suite 30 min prior to the start of the euthanasia protocol (9:00 or 11:00, respective to fast length), and were euthanized one at a time. Weight gain was calculated as the difference between the final body weight of the rats following the fast and the body weight at diet start. Trunk blood was collected and serum saved and stored at -80 °C. Body length from nose to anus and tail length from anus to tail tip were measured with a ruler. The brain, visceral fat pad tissues (retroperitoneal (RetroFat), omental/mesenteric fat (OmenFat), and parametrial fat (ParaFat)), liver, kidneys, and heart were dissected, weighed, and snap-frozen. Sections of the ileum and colon were also dissected and snap-frozen.

WKY/NCrl tissues were collected for qPCR analyses, and rats were euthanized via decapitation after no fast. Rats were transferred to the anteroom of the necropsy suite 30 min prior to the start of euthanasia (9:00) and were euthanized one at a time. The brain, visceral fat pad tissues, liver, kidneys, and heart were dissected and snap-frozen in both male and female rats. We also collected pituitary, adrenals, sections of the ileum and colon, subcutaneous white and brown adipose tissue, and soleus muscle from male inbred rats, but based on findings in male rats, additional tissues collected in the females included the pituitary, adrenals, ileum, and colon.

### Insulin

We used ultrasensitive ELISA kits (Alpco Ref # 80-INSTRU-E10) to analyze fasting serum insulin collected from the IPGTT.

### Corticosterone

We used a corticosterone (CORT) competitive ELISA kit (ThermoFisher Ref # EIACORT) to analyze serum corticosterone from trunk blood collected at euthanasia. Samples from HFD females in Study 1 were sent to the Redei lab at Northwestern University, while all other samples were run in-house at WFSoM. As guided by kit instructions, samples were diluted at least 1:100, and analyzed at 450 nm against a 4-parameter standard curve. The ELISA kit is confirmed to preferentially bind the active corticosterone molecule.

### RNA Extraction

For both experimental animals and inbred animals, RNA was extracted from fatty tissues using the RNeasy Lipid Tissue Mini Kit (Qiagen Cat # 74804). Non-fatty tissue, such as liver or intestine, were extracted by Trizol.

### Real-time quantitative PCR

We assessed expression of *Krtcap3* (liver) and *adenylate cyclase 3* (*Adcy3*) (RetroFat; **Table 1**) between Study 1 and Study 2 rats to determine if there had been changes in gene expression that could contribute to the change in phenotype we saw. We examined *Krtcap3* expression in the liver only in WT animals but examined *Adcy3* expression in the RetroFat for both studies and genotypes. We chose to investigate expression of *Adcy3* in addition to *Krtcap3* because of its well-known influence on adiposity (14, 15) and the close proximity of the genes to each other (10).

**Table 1.**
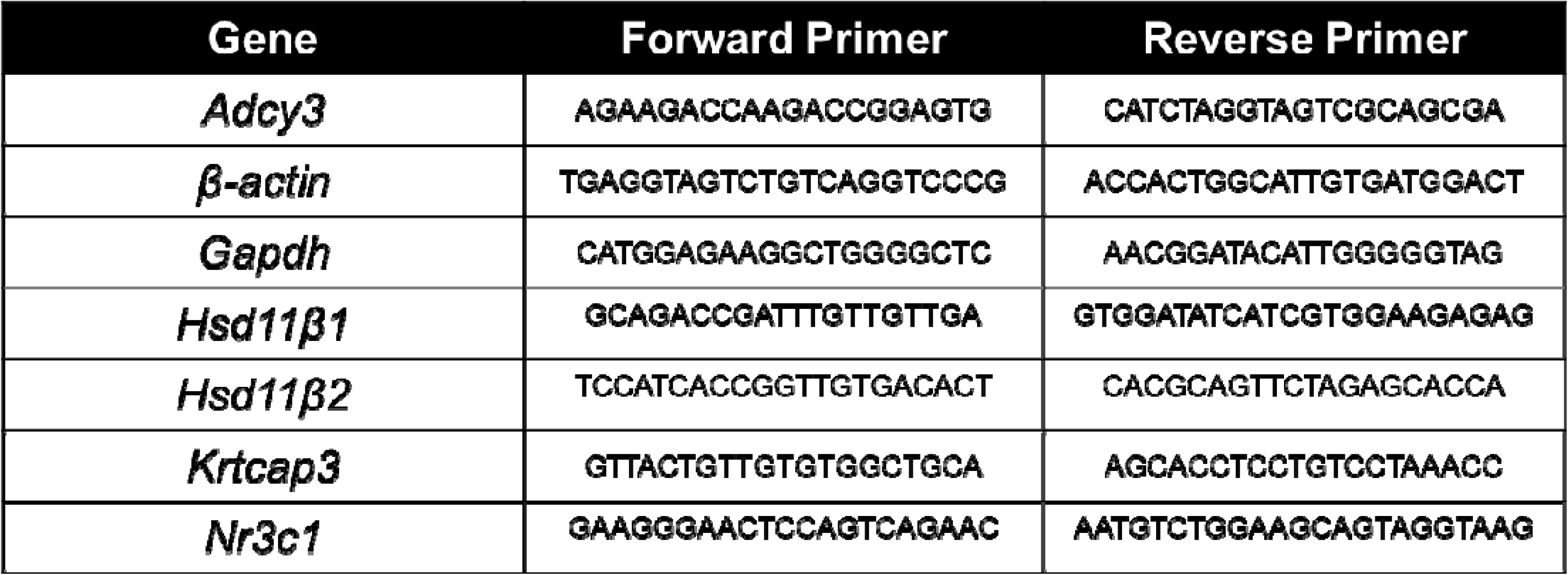
Primer sequences. 5’ ➔ 3’

We examined expression of several stress-related genes in liver and RetroFat between Study 1 and Study 2 WT and KO to evaluate if genotype or environment affected gene expression. Liver and fat were selected because both tissues had been collected in both studies. We analyzed *Nr3c1, Hsd11*β*1,* and *Hsd11*β*2* (**Table 1**) in both sets of tissues. Nuclear receptor subfamily 3 group C member 1 (NR3C1) is a glucocorticoid (GC) receptor, and the 11β-hydroxysteroid dehydrogenases (11β-HSDs) are responsible for catalyzing corticosterone, where isoform 1 (11β-HSD1) primarily activates the pathway and isoform 2 (11β-HSD2) de-activates the pathway.

To measure gene expression, either *Gapdh* or β*-actin* were used as housekeeping genes (**Table 1**). Fold change of the corresponding gene of interest transcript was calculated by the following equation:

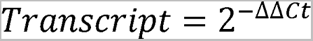

Where ΔCt was the difference between the crossing threshold (Ct) of the gene of interest and the housekeeping gene, and ΔΔCt the difference between each sample ΔCt and the average control ΔCt.

To measure gene expression in the diet study rats, the average control ΔCt was the S1 WT mean ΔCt values, respective to each gene and tissue. To measure *Krtcap3* expression in WKY/NCrl rats, expression of *Krtcap3* in the WKY male pituitary was used as the average control ΔCt.

### RNA-seq

Sections of hypothalamus were dissected from frozen brain of select female rats from Study 1 and Study 2 (n = 4/study and genotype). Although *Krtcap3* has low expression in the hypothalamus, the hypothalamus is well-connected to satiety signaling (16) and the hypothalamic-pituitary-adrenal (HPA) axis of stress response (17), and we aimed to identify changes in these pathways. We chose rats that were the first rat of their cage to be euthanized and were representative of phenotype differences between WT and KO rats. Brains had been snap-frozen at euthanasia and stored at -80C, were dissected at -20C, and then the tissue stored again at -80C until they were shipped to Azenta (Genomics from Azenta Life Sciences, South Plainfield, NJ).

#### RNA Extraction

Total RNA was extracted from fresh frozen tissue samples using Qiagen RNeasy Plus Universal mini kit (Qiagen, Hilden, Germany) following manufacturer’s instructions.

#### Library Preparation with PolyA Selection

RNA samples were quantified using Qubit 2.0 Fluorometer (Life Technologies, Carlsbad, CA, USA) and RNA integrity was checked using Agilent TapeStation 4200 (Agilent Technologies, Palo Alto, CA, USA). For all samples, the RNA concentration was at least 200 ng/uL, and the RIN was at least 7.9.

RNA sequencing libraries were prepared using the NEBNext Ultra II RNA Library Prep for Illumina using manufacturer’s instructions (NEB, Ipswich, MA, USA). Briefly, mRNAs were initially enriched with Oligod(T) beads. Enriched mRNAs were fragmented for 15 minutes at 94 °C. First strand and second strand cDNA were subsequently synthesized. cDNA fragments were end repaired and adenylated at 3’ends, and universal adapters were ligated to cDNA fragments, followed by index addition and library enrichment by PCR with limited cycles. The sequencing libraries were validated on the Agilent TapeStation, and quantified by using Qubit 2.0 Fluorometer as well as by quantitative PCR (KAPA Biosystems, Wilmington, MA, USA).

#### Illumina Sequencing

The sequencing libraries were clustered on a lane of a HiSeq flowcell. After clustering, the flowcell was loaded on the Illumina instrument (4000 or equivalent) according to manufacturer’s instructions. The samples were sequenced using a 2x150bp Paired End (PE) configuration. Image analysis and base calling were conducted by the Control software. Raw sequence data (.bcl files) generated the sequencer were converted into fastq files and de-multiplexed using Illumina’s bcl2fastq 2.17 software. One mismatch was allowed for index sequence identification.

#### Differentially Expressed Gene Analysis

After investigating the quality of the raw data, sequence reads were trimmed to remove possible adapter sequences and nucleotides with poor quality using Trimmomatic v.0.36. The trimmed reads were mapped to the Rattus norvegicus reference genome RN6 available on ENSEMBL using the STAR aligner v.2.5.2b. BAM files were generated as a result of this step. Unique gene hit counts were calculated by using feature Counts from the Subread package v.1.5.2. Only unique reads that fell within exon regions were counted. After extraction of gene hit counts and normalization, the gene hit counts table was used for downstream differential expression analysis. Using DESeq2, comparisons of gene expression between the groups of samples (WT vs KO and Study 1 vs Study 2) were performed (**Table 2**). The Wald test was used to generate p values and log2 fold changes. In our analysis, genes with p values < 0.05 were called as significantly differentially expressed genes for each comparison.

**Table 2.** Comparison groups for differentially-expressed genes in the RNA-seq analysis. We analyzed differences between wild-type (WT) and *Krtcap3* knock-out (KO) for each study, and then differences by study for each genotype.

|  | Group 1 | Group 2 |
| --- | --- | --- |
| Comparison 1 | Study 1 WT | Study 1 KO |
| Comparison 2 | Study 2 WT | Study 2 KO |
| Comparison 3 | Study 1 WT | Study 2 WT |
| Comparison 4 | Study 1 KO | Study 2 KO |

### Ingenuity Pathway Analysis (IPA)

We used IPA software (QIAGEN Digital Insights) to analyze differentially expressed genes in the context of known biological responses and regulatory networks. Differentially expressed genes in each comparison were uploaded to the IPA software. We identified canonical pathways that differed between each comparison and networks that included both direct and indirect relationships, with significance determined in IPA by Fisher’s Exact Test and the threshold set at –log(p-value) > 1.3. We then mapped upstream regulators that were activated with a p-value of overlap < 0.05, and removed indirect relationships as well as orphan nodes.

### Statistical Analysis

All data were analyzed in R (1.4.1103), with the exception of RNA-seq data, which was analyzed as described above, and cumulative energy intake and WKY/NCrl tissue expression data which were analyzed in GraphPad PRISM (9.4.1(681)). Outliers according to Grubb’s test were removed, distribution was assessed by the Shapiro-Wilks test and data were transformed to reflect a normal distribution if necessary. Homogeneity of variance was assessed by Levene’s test.

In examining data collected from Study 2, we first analyzed the effect of genotype. All single point adiposity measurements were analyzed by Student’s t-test. Growth, food/energy consumption, and energy expenditure curves were assessed with a repeated measures ANOVA, where the effect of genotype was examined over time.

When analyzing phenotypes between Study 1 and Study 2, most of the metabolic data were analyzed by a two-way ANOVA, where the factors were genotype and study. Please see **Appendix Table 1** for adiposity measures from Study 1 rats. Growth and food consumption curves were analyzed by a two-way repeated measures ANOVA, examining the effects of genotype and study over time. If there was a significant interaction, the analysis was split by genotype and either a Student’s t-test or a repeated measures ANOVA (respective to the phenotype) assessed the effect of study. Slopes of the cumulative energy intake curves were tested in PRISM to determine if they were significantly different, and then split by genotype to examine slopes between studies.

When examining the results from the CORT assays, we noticed the data was spread in an almost bimodal fashion, particularly for KO animals. Upon further investigation, the factor that most likely explained these differences was euthanasia order within the cage—which rat had been euthanized first and which rat had been euthanized second. Due to this, we included euthanasia order as a third factor in the analyses, in addition to genotype and study. The data were analyzed by a 3-way ANOVA, and split according to interactions that were significant.

In the data comparing male and female *Krtcap3* expression in multiple tissues from naïve WKY/NCrl rats, male and female data were compared for each tissue individually using multiple comparisons t-tests corrected with the Holm-Šidák test.

## Results

### No difference in adiposity measures between WT and KO rats

In contrast to our previous findings (13), there were no differences in body weight between WT and KO females, neither at wean nor at six weeks of age, nor were there differences in adiposity (**Table 3**). Unexpectedly, no differences ever developed in the weight between WT and KO females (**Figure 2a**), nor in food consumption (**Figure 2b**). During the week of individual housing WT and KO females consumed similar amounts of calories per day (**Table 3**). We ultimately did not see any differences in weight gain between WT and KO females (**Table 3**), and at euthanasia KO females were not heavier than WT females (**Table 3**).

**Figure 2.**
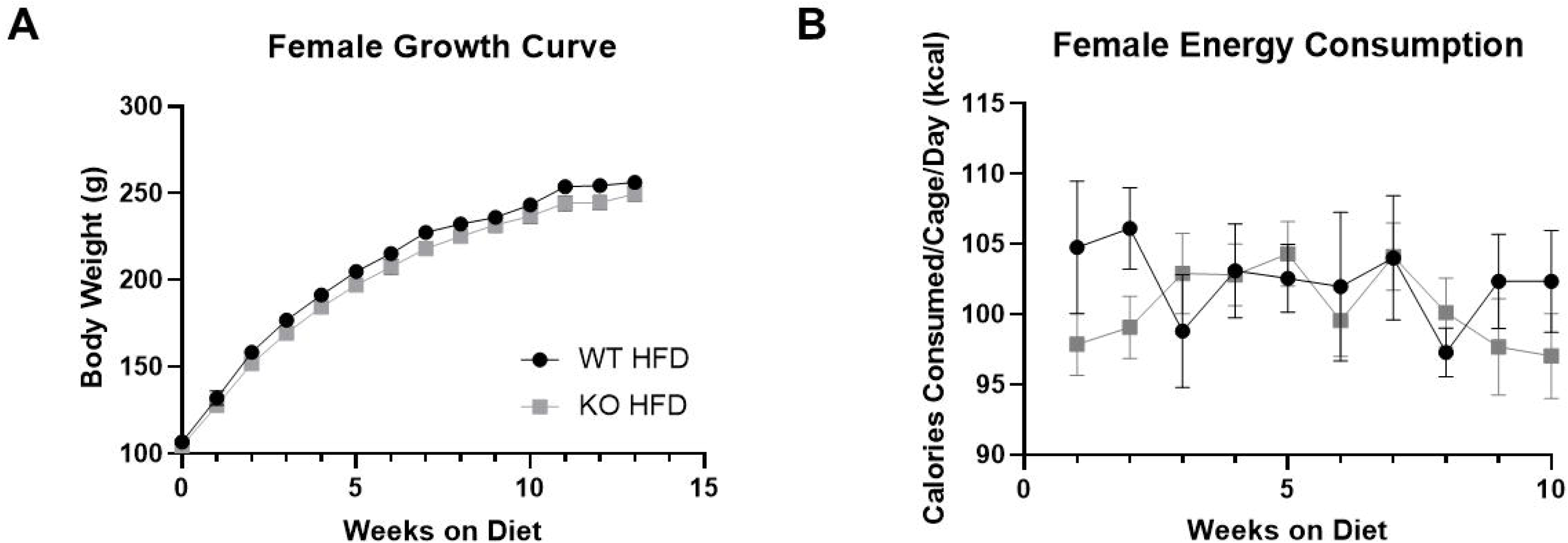
Body weight growth and food consumption in wild-type (WT) and *Krtcap3* knock-out (KO) rats. Rats were weighed weekly throughout the study, and food consumption per cage was recorded for the first 10 weeks on diet. (A) There were no significant differences between the growth curve of WT (black circle) and KO (gray square) female rats. (B) There were also no differences in the food consumption patterns between WT and KO cages over the course of the study.

**Table 3.**
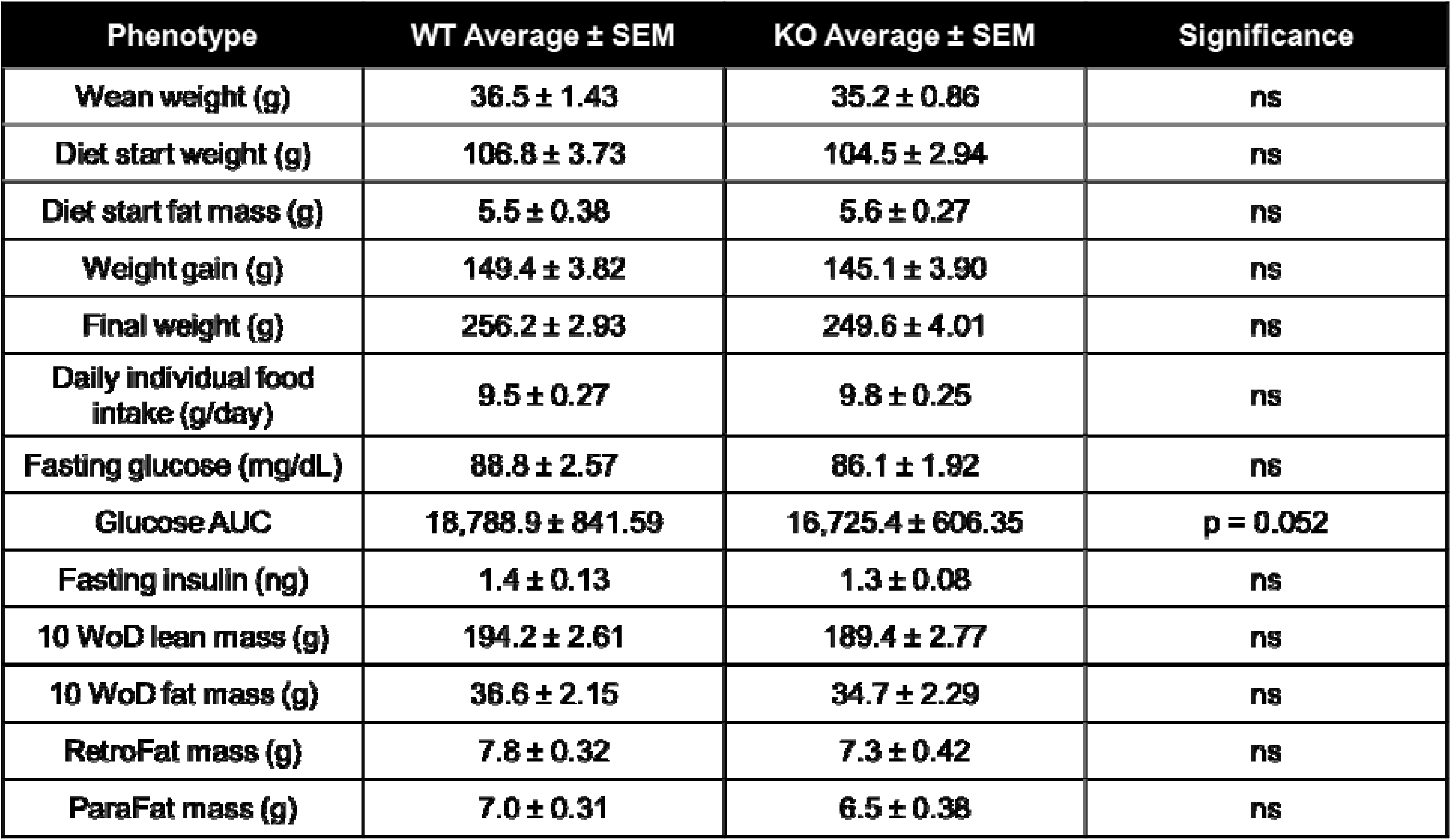
Adiposity measures between wild-type (WT) and *Krtcap3* knock-out (KO) female rats from Study 2. Weeks on diet (WoD), area under the curve (AUC), retroperitoneal fat (RetroFat), and parametrial fat (ParaFat). Values are given as the mean average ± standard error. Significance is indicated in the final column.

| Phenotype | WT Average ± SEM | KO Average ± SEM | Significance |
| --- | --- | --- | --- |
| Wean weight (g) | 36.5 ± 1.43 | 35.2 ± 0.86 | ns |
| Diet start weight (g) | 106.8 ± 3.73 | 104.5 ± 2.94 | ns |
| Diet start fat mass (g) | 5.5 ± 0.38 | 5.6 ± 0.27 | ns |
| Weight gain (g) | 149.4 ± 3.82 | 145.1 ± 3.90 | ns |
| Final weight (g) | 256.2 ± 2.93 | 249.6 ± 4.01 | ns |
| Daily individual food intake (g/day) | 9.5 ± 0.27 | 9.8 ± 0.25 | ns |
| Fasting glucose (mg/dL) | 88.8 ± 2.57 | 86.1 ± 1.92 | ns |
| Glucose AUC | 18,788.9 ± 841.59 | 16,725.4 ± 606.35 | p = 0.052 |
| Fasting insulin (ng) | 1.4 ± 0.13 | 1.3 ± 0.08 | ns |
| 10 WoD lean mass (g) | 194.2 ± 2.61 | 189.4 ± 2.77 | ns |
| 10 WoD fat mass (g) | 36.6 ± 2.15 | 34.7 ± 2.29 | ns |
| RetroFat mass (g) | 7.8 ± 0.32 | 7.3 ± 0.42 | ns |
| ParaFat mass (g) | 7.0 ± 0.31 | 6.5 ± 0.38 | ns |

WT and KO females had comparable fasting glucose levels (**Table 3**) though unlike the prior study, WT females did have a slightly increased glucose AUC compared to KO females (T_27_ = 2.03, p = 0.052; **Table 3**). Surprisingly, there were no differences in fasting insulin between WT and KO females (**Table 3**).

Over the 24 h measurement period there were no differences in total energy expenditure between WT and KO rats when normalized to body weight (**Appendix Figure 1a**). There were also no differences in respiratory exchange ratio between the genotypes (**Appendix Figure 1b**), with comparable usages of fats and carbohydrates as fuel sources. Finally, while a visual difference develops in food intake between WT and KO rats (**Appendix Figure 1c**), this difference is ultimately not significant, corroborating the other measures of food intake throughout the study.

At EchoMRI analysis after 10 weeks on diet, WT and KO females had similar amounts of total lean mass and fat mass (**Table 3**). At euthanasia, there were no differences in RetroFat mass nor ParaFat mass between WT and KO females (**Table 3**). Though these data contradict our previous findings (13), the adiposity and metabolic measures do support the lack of body weight differences seen between WT and KO females in this study. There were also no noteworthy differences in body size or non-adipose organ weights between WT and KO rats (**Table 4**).

**Table 4.** Body size and non-adipose organ weights between wild-type (WT) and *Krtcap3* knock-out (KO) female rats. Body length was measured from nose to anus, and tail length was measured from anus to end of tail. Values are given as the mean average ± standard error. Significance is indicated in the final column.

| Phenotype | WT Average ± SEM | KO Average ± SEM | Significance |
| --- | --- | --- | --- |
| Body Length (cm) | 21.4 ± 0.13 | 21.2 ± 0.12 | ns |
| Tail Length (cm) | 16.9 ± 0.19 | 16.4 ± 0.6 | p = 0.09 |
| Brain Weight (g) | 1.9 ± 0.02 | 1.9 ± 0.02 | ns |
| Liver Weight (g) | 7.6 ± 0.11 | 7.7 ± 0.16 | ns |
| Heart Weight (g) | 1.1 ± 0.09 | 1.0 ± 0.03 | p = 0.092 |
| Kidney Weight (g) | 0.8 ± 0.02 | 0.7 ± 0.01 | ns |

### WT rats in Study 2 are heavier and eating more than in Study 1, with no change in KO

When examining differences between Study 1 and Study 2, we found no differences in body weight at study start between Study 1 and Study 2 females of either genotype (**Figure 3a**). However, when we examined body weight over time, we saw a significant interaction between genotype and study (**Figure 3b**; F_1, 44_ = 10.3, p = 0.002) where Study 2 WT females were heavier over time compared to Study 1 WT females (F_2.46, 44.22_ = 8.02, p = 5.05e-4), with no effect in KO females.

**Figure 3.**
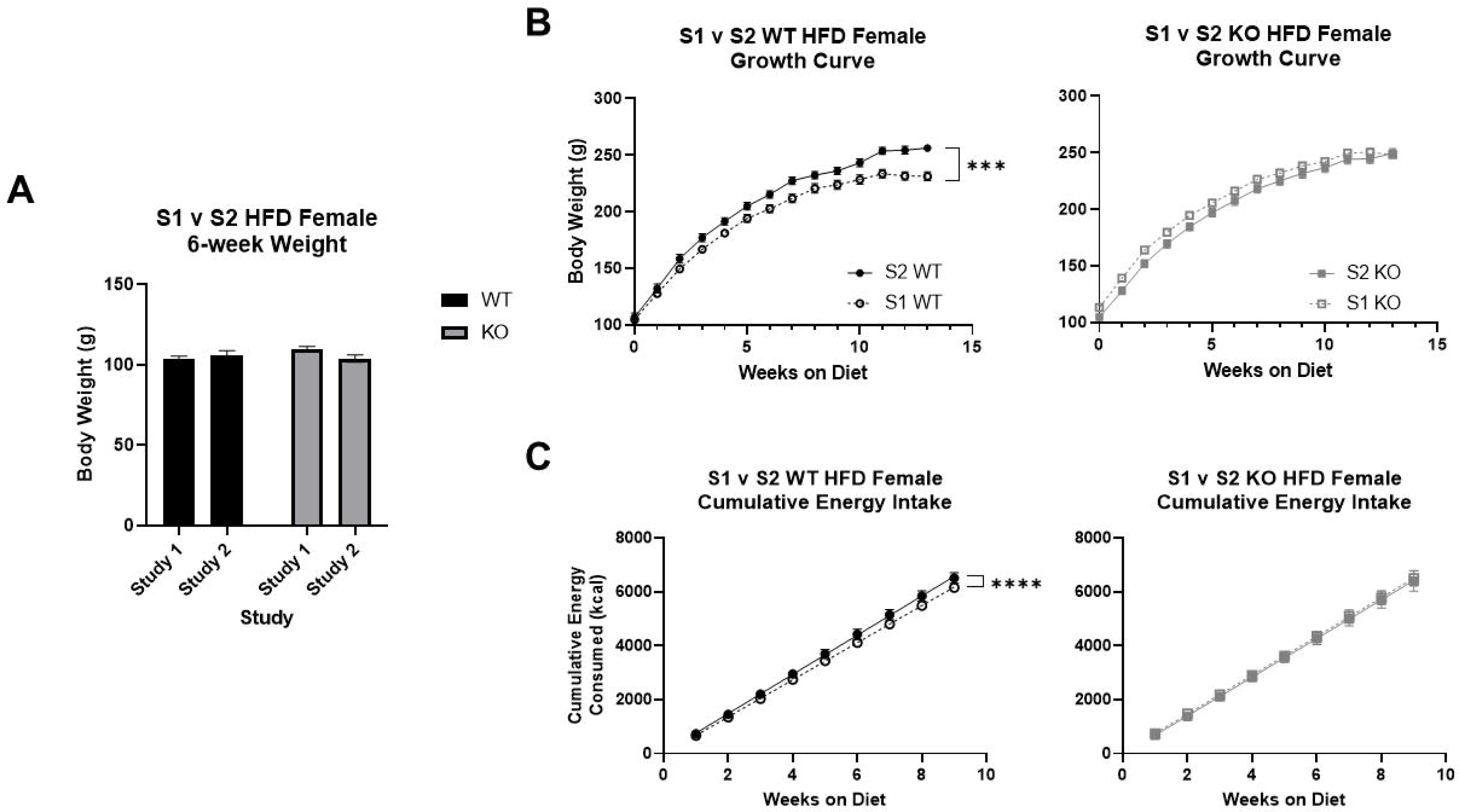
Differences in select adiposity phenotypes between Study 1 (S1) and Study 2 (S2) in wild-type (WT) and *Krtcap3* knock-out (KO) rats. (A) Comparing body weight at diet start, or six weeks of age, between the two studies, there are no differences in the starting weight of the WT rats (black) or the KO rats (gray). However, (B) Study 2 WT rats (filled circle, solid line) gained significantly more weight over the course of the study compared to Study 1 WT rats (open circle, dashed line). There were no comparable differences between the growth curves of Study 1 KO rats (open square, dashed line) and Study 2 KO rats (filled square, solid line). Perhaps explaining these differences in body weight curves, (C) Study 2 WT rats had a greater cumulative energy consumption compared to Study 1 WT rats, while there were no differences between the KO rats. ***p < 0.001, ****p < 0.0001 represent the effect of study and weeks on diet respective to each genotype.

We then compared food consumption between the two studies. There was a significant interaction between study and genotype (F_1, 17_ = 4.55, p = 0.048). WT rats in Study 2 consumed significantly more food than WT rats from Study 1 (F_1, 7_ = 6.72, p = 0.036; **Appendix Figure 2**), with no changes between studies in KO rats. When confirming caloric intake over the study, there was a near interaction between genotype and study (F_1, 19_ = 3.24, p = 0.088) and a main effect of week (F_9, 171_ = 3.36, p = 8.14e-4). To better examine the difference in energy intake over time between study and genotype, we calculated cumulative energy intake and compared the slopes of the curves. While Study 2 WT females had a greater caloric intake than Study 1 WT females (F_1, 198_ = 6.94e14, p < 0.0001; **Figure 3c**), there were no changes between studies in KO caloric intake.

We next compared different adiposity measures between Study 1 and Study 2 in WT and KO rats, namely the measurements recorded by EchoMRI analysis and those collected at euthanasia. At EchoMRI analysis after 10 weeks on diet, the spread of the data for total fat mass was too large to see statistically significant effects, but there is a visual increase in Study 2 WT rats compared to Study 1 that is not present inthe KO rats (**Figure 4a**). There was, however, a significant interaction between genotype and study for total lean mass (F_1, 45_ = 5.96, p = 0.019) where Study 2 WT rats had significantly greater lean mass compared to Study 1 counterparts (T_19_ = 2.20, p = 0.041; **Figure 4b**), with no difference in KO rats.

**Figure 4.**
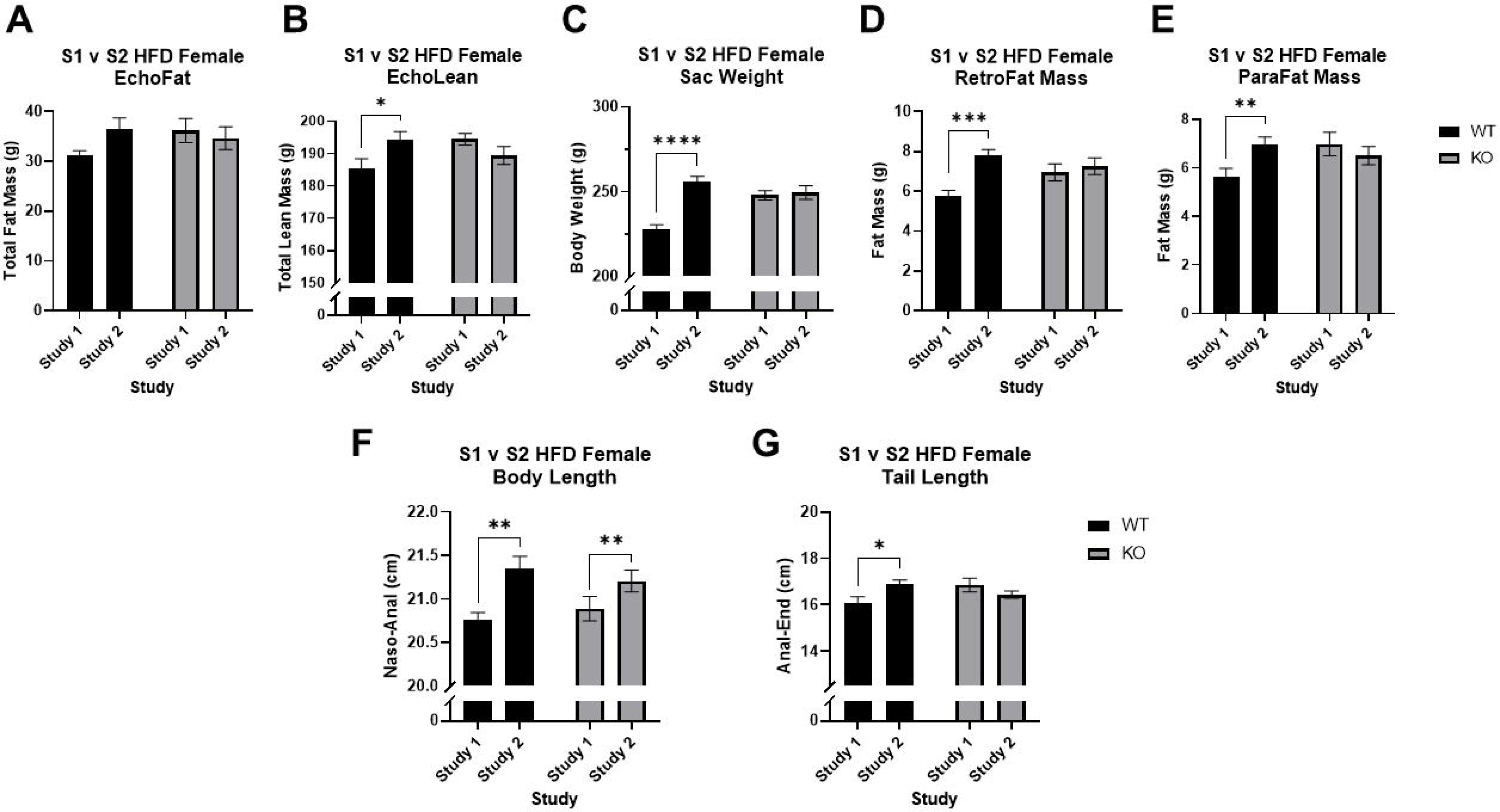
Differences in size between Study 1 (S1) and Study 2 (S2) wild-type (WT) and *Krtcap3* knock-out (KO) rats. Total fat mass (A) and total lean mass (B) were recorded after 10 weeks on diet. At euthanasia three weeks later body weight (C) plus retroperitoneal fat (RetroFat; D) and parametrial fat (ParaFat; E) mass were recorded. Body length measured from nose to anus (F) and tail length measured from anus to end of tail (G) were also recorded at this time. (A) There were no differences in total fat mass by study in either WT (black) or KO (gray) rats at EchoMRI analysis, though there was a visual increase in S2 WT rats. (B) S2 WT rats did have a greater total lean mass compared to S1 counterparts, with no differences in KO rats. (C) In WT rats, Study 2 rats had an increased final body weight compared to Study 1 rats, with no changes between the final body weights of KO rats. (D-E) Study 2 WT rats had greater RetroFat mass and ParaFat mass compared to Study 1 WT rats, while there were no significant differences in the KO rats. (F) There was a significant increase in the body length of Study 2 WT and KO rats but (G) there was only a significant increase in tail length in Study 2 WT animals and no difference in KO rats. *p < 0.05, **p < 0.01, ***p < 0.001, ****p < 0.0001 represent the effect of study respective to each genotype, with the exception of the body length phenotype, which represents the main effect of study.

The adiposity differences were more prominent three weeks later at euthanasia, likely because WT rats in Study 2 continued to grow in size while WT rats in Study 1 appeared to plateau (**Figure 3b**). Specifically, there was a significant interaction between study and genotype (F_1, 44_ = 12.5, p = 9.72e-4) where Study 2 WT rats were much heavier than Study 1 WT rats (T_18_ = 7.19, p = 1.09e-6; **Figure 4c**), with no differences in KO rats. Similar trends were seen in the measures for visceral fat mass: there was an interaction between study and genotype for RetroFat mass (F_1, 45_ = 4.34, p = 0.043) where Study 2 WT females had a significant increase in RetroFat mass compared to Study 1 WT females (T_19_ = 4.71, p = 1.5e-4; **Figure 4d**), but no differences in KO females. This was also true for ParaFat mass: there was an interaction between study and genotype (F_1, 45_ = 4.92, p = 0.032) but ultimately the only difference was in WT rats (T_19_ = 2.89, p = 9.3e-3; **Figure 4e**). Interestingly, when examining the data from EchoMRI analysis normalized to body weight, there were no differences in total body fat percentage nor in total body lean percentage (data not shown).

There was a statistically significant increase in the body length of Study 2 rats compared to Study 1 rats for both genotypes (F_1, 44_ = 11.6, p = 0.001; **Figure 4f**). There was also a significant interaction between study and genotype for tail length (F_1, 45_ = 7.4, p = 0.009), where only Study 2 WT rats had a longer tail length compared to Study 1 (T_19_ = 2.53, p = 0.021; **Figure 4g**). There were no statistical differences in brain weight (**Appendix Figure 3a**) nor in heart weight (**Appendix Figure 3b**) between the studies, but Study 2 rats had an increased average kidney weight compared to Study 1 rats for both genotypes (F_1, 45_ = 8.76, p = 0.005; **Appendix Figure 3c**).

### No changes in Krtcap3 nor Adcy3 expression

We measured liver *Krtcap3* between Study 1 and Study 2 WT rats to confirm expression had not diminished, and found no differences (**Appendix Figure 4a)**. We did not include KO rats in this analysis as we previously demonstrated KO rats had significantly reduced *Krtcap3* expression in the liver (13). We also measured RetroFat *Adcy3* expression to verify its expression was not interfering with the phenotype, but found no differences by genotype nor by study (**Appendix Figure 4b**).

### Differences in basal and stress CORT between WT and KO rats is dependent on study and cage order

When we initially assessed CORT collected from trunk blood at euthanasia, there were no significant differences by either genotype or study (**Figure 5a**), though there was a slight trend towards an interaction between study and genotype (F_1, 38_ = 2.95, p = 0.094). Closer investigation of the data revealed a bimodal pattern to the data that we determined was related to the order of euthanasia in the cage. When we accounted for order as a factor, there were interactions between study and genotype (F_1, 34_ = 5.49, p = 0.025) and between genotype and order (F_1, 34_ = 12.68, p = 0.001).

**Figure 5.**
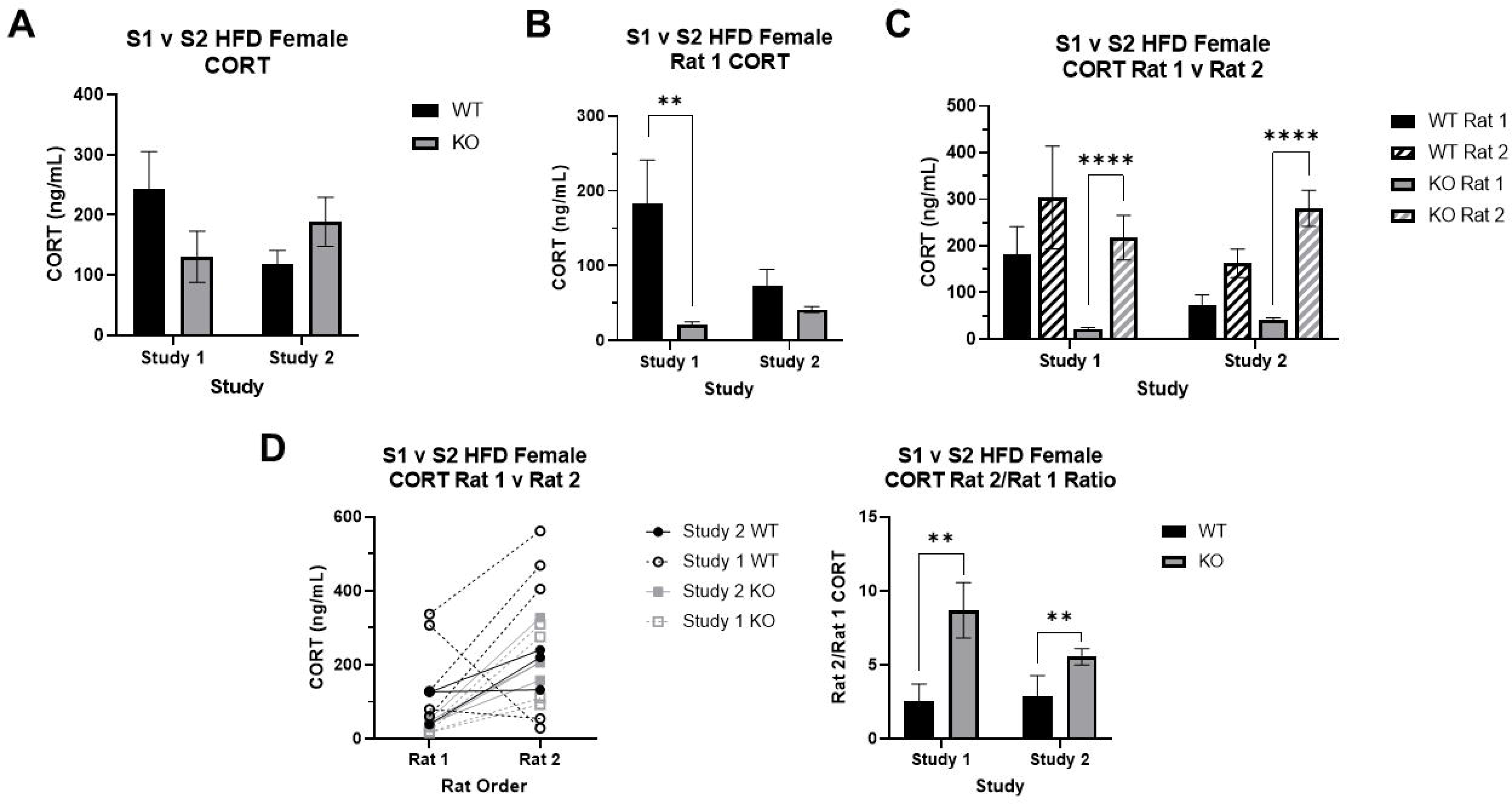
Corticosterone (CORT) measurements between Study 1 (S1) and Study 2 (S2) wild-type (WT) and *Krtcap3* knock-out (KO) rats. CORT was measured in the trunk blood collected at euthanasia. (A) All rats evaluated together. When grouped only by study and genotype, there are no significant differences between WT (black) and KO (gray) CORT in either Study 1 or Study 2. Further assessment of the data revealed a third factor to consider, the order within the cage the rats were euthanized, either first (Rat 1) or second (Rat 2). Re-analysis with this additional variable revealed a significant interaction between study and genotype as well as one between genotype and order. (B) Assessing only rats euthanized first (basal CORT). KO rats had lower CORT than WT rats in both studies, but this was statistically significant only in Study 1, not Study 2. **p < 0.01 represents the effect of genotype respective to each study. (C) Comparing between rats euthanized first and second. In both studies, KO CORT significantly increases in Rat 2 of the cage, an effect that is not see in the WT rats. ****p < 0.0001 represents the effect of order respective to each genotype. (D) Rat 2/Rat 1 CORT ratio. There was a significant main effect of genotype, where KO rats of both studies had a greater ratio than WT rats, indicating a greater CORT response. **p < 0.01 represents a main effect of genotype.

We first split the data by euthanasia order—analyzing rats euthanized first (Rat 1) separately from those euthanized second (Rat 2). For rats euthanized first, what we consider to be a true basal measure, there was a main effect of genotype (F_1, 15_ = 21.2, p = 3.4e-4) where KO rats had lower CORT than WT rats (**Figure 5b**). There was also an interaction between study and genotype (F_1, 15_ = 8.69, p = 0.01). Specifically, WT rats from Study 1 showed significantly higher basal CORT relative to KO rats (T_5.21_ = 5.16, p = 3.17e-3; **Figure 5b**), with no significant difference by genotype in Study 2. Additionally, WT rats in Study 2 had lower CORT than those in Study 1 (T_7.79_ = 1.88, p = 0.098); while CORT in KO rats did increase between the two studies (T_5.99_ = 3.71, p = 0.01), the difference is likely not physiologically meaningful. There were no differences by study nor by genotype in the rats euthanized second.

We then included both Rat 1 and Rat 2 and analyzed the data by genotype to assess the effect of study and order. We found that study (F_1, 18_ = 8.08, p = 0.011) and euthanasia order (F_1, 18_ = 143.15, p = 5.29e-10; **Figure 5c**) strongly impacted CORT in the KO rats, but had no effect in the WT rats. Specifically, Study 2 KO rats had a slightly higher basal CORT compared to Study 1 counterparts, while in both studies there was a major CORT increase in the KO rats that were euthanized second.

Finally, to continue evaluating the differences in CORT response between WT and KO rats at euthanasia, we plotted the change in CORT between Rat 1 and Rat 2 for each cage, and then calculated the ratio increase of Rat 2 CORT compared to Rat 1. There was a main effect of genotype, where KO rats of both studies had a significantly higher CORT ratio compared to WT rats (F_1, 14_ = 12.58, p = 0.003; **Figure 5d**). These data demonstrate a stronger acute social stress response in KO relative to WT rats, specifically in Rat 2’s response to the removal of Rat 1.

### Changes in expression of genes associated with glucocorticoid processing indicate different stress environments between Study 1 and Study 2

We measured expression of genes associated with CORT function and metabolism in liver and fat from Study 1 and Study 2 rats to evaluate if expression was different by genotype or by study.

In liver tissue, there was higher expression of the glucocorticoid receptor *Nr3c1* in Study 1 compared to Study 2 (F_1, 22_ = 65.1, p = 7.18e-8; **Figure 6a**) but no difference by genotype in either study. Expression of the hydroxysteroid dehydrogenases followed a similar pattern, with expression elevated in Study 1 for both *Hsd11*β*1* (F_1, 22_ = 19.35, p = 2.3e-4; **Figure 6b**) and *Hsd11*β*2* (F_1, 20_ = 24.35, p = 97.99e-5; **Figure 6c**), though ultimately no genotype differences in either.

**Figure 6.**
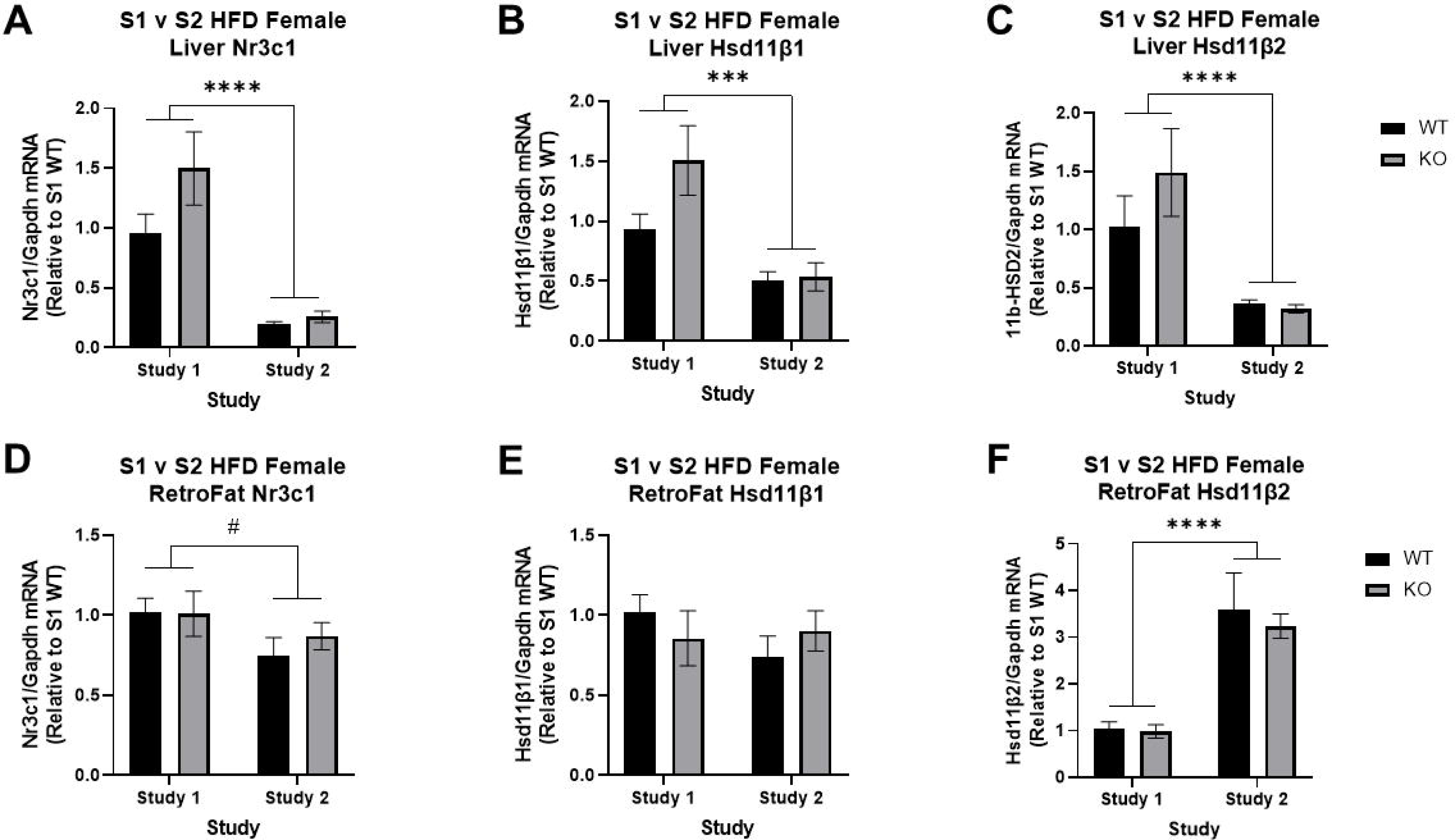
Expression of the glucocorticoid receptor and genes associated with corticosteroid processing between Study 1 (S1) and Study 2 (S2) in wild-type (WT) and *Krtcap3* knock-out (KO) rats. We measured expression of *nuclear receptor subfamily 3 group C member 1* (*Nr3c1*), *11*β*-hydroxysteroid dehydrogenase isoform 1* (*Hsd11*β*1*) and *11*β*-hydroxysteroid dehydrogenase isoform 2* (*Hsd11*β*2*) in liver and retroperitoneal fat (RetroFat) tissue of rats from both studies. In the liver, (A) Study 1 WT (black) and KO (gray) rats had significantly higher expression of *NR3C1* compared to Study 2 rats, with no differences by genotype. Similarly, Study 1 rats of both genotypes had higher expression of (B) *Hsd11*β*1* and (C) *Hsd11*β*2* when compared to Study 2 rats. In the RetroFat, (D) Study 1 WT and KO rats had slightly higher *Nr3c1* when compared to Study 2 rats. (E) There were no differences in *Hsd11*β*1* expression between the two studies, but (F) Study 2 rats had significantly higher *Hsd11*β*2* compared to Study 1 rats. #p < 0.1, ***p < 0.001, ****p < 0.0001 represent a main effect of study.

In RetroFat tissue *Nr3c1* expression was also slightly higher in Study 1 compared to Study 2 (F_1, 18_ = 3.54, p = 0.076; **Figure 6d**), though still no differences by genotype. The hydroxysteroid dehydrogenases, on the other hand, displayed different patterns in RetroFat tissue compared to liver. While there were no differences in *Hsd11*β*1* expression by study nor by genotype (**Figure 6e**), *Hsd11*β*2* expression was lower in Study 1 compared to Study 2 (F_1, 17_ = 47.55, p = 2.59e-6; **Figure 6f**).

### Krtcap3 is highly expressed in both sexes along the gastrointestinal tract and in the pituitary gland, with differences in expression between males and females in the gonads and pituitary

Using naïve WKY/NCrl male rats, *Krtcap3* had low expression in all of the different fat pads, the adrenals, the heart, the muscle, and different regions of the hypothalamus (**Figure 7a**). Expression between the liver and kidney was comparable while expression in the pituitary, testes, and gastrointestinal tract was high.

**Figure 7.**
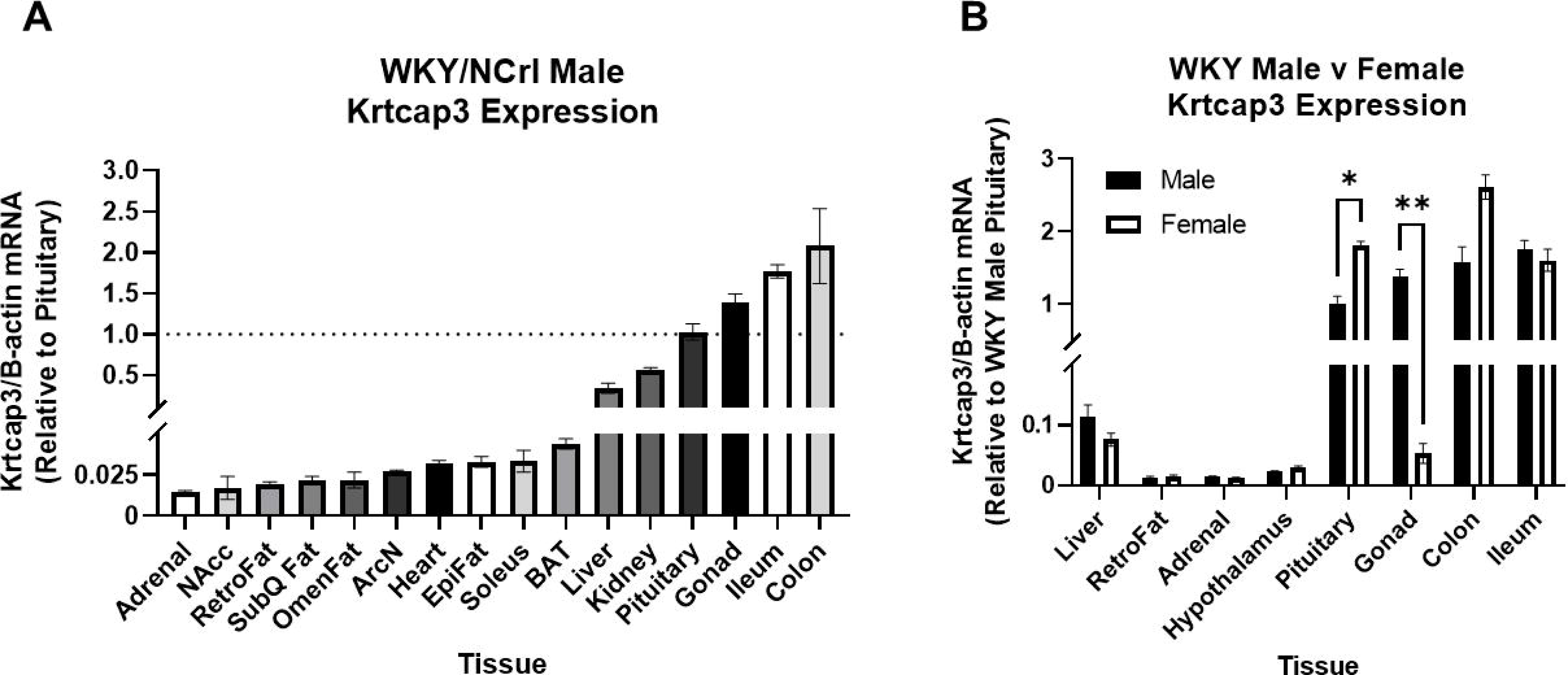
*Krtcap3* expression in multiple tissues between male and female naïve Wistar-Kyoto (WKY/NCrl) rats ordered from Charles-River and euthanized for tissue collection. Tissues include: liver, adrenal, nucleus accumbens (NAcc), retroperitoneal fat (RetroFat), subcutaneous white fat (SubQ Fat), omental/mesenteric fat (OmenFat), arcuate nucleus (ArcN), heart, Epididymal fat (EpiFat), soleus muscle, subcutaneous brown fat (BAT), kidney, pituitary, gonads (testes or ovaries, respectively), whole hypothalamus, ileum, and colon. (A) Comparing *Krtcap3* expression in male WKY rats across multiple tissues, nearly all tissues had lower expression when compared to pituitary *Krtcap3* expression, particularly the adrenal gland, different brain regions, and liver. Noteworthy tissues with higher *Krtcap3* expression than pituitary are the male gonad and the intestines. The dotted line across the plot represents a fold change of one, or *Krtcap3* expression in the pituitary. (B) When comparing *Krtcap3* expression between male and female WKY rats in select tissues, there were significant sex differences in two tissues: the pituitary where females had higher *Krtcap3* expression than males and the gonads where females had lower *Krtcap3* expression than males. For inter-tissue visual comparison, all tissues were normalized to *Krtcap3* expression in the WKY male pituitary. *p < 0.05, **p < 0.01 for sex differences, adjusted for multiple comparisons.

We then assessed *Krtcap3* expression in select tissues between naïve male and female WKY rats to establish any differences in expression by sex (**Figure 7b**). Female rats had significantly higher *Krtcap3* expression in the pituitary (T_4_ = 7.05, p = 0.015) compared to male rats. Male rats, on the other hand, had much higher *Krtcap3* expression in the testes than females did the ovaries (T_4_ = 13.41, p = 1.43e-3). There was an additional visual difference between the sexes for *Krtcap3* expression in the colon, but it did not meet the significance threshold.

### Pathway analysis of hypothalamic RNA-seq data indicate improved myelination

We assessed changes in gene expression in the hypothalamus of WT and KO rats of both Study 1 and Study 2. Although we have shown that *Krtcap3* has very low expression in the hypothalamus, we aimed to identify changes in either satiety signaling or stress response signaling pathways that could indicate which pathways *Krtcap3* may be involved in. We identified 132 significant differentially expressed genes (DEGs) between Study 1 WT and KO female rats (**Table S1**) and 290 significant DEGs between Study 2 WT and KO rats (**Table S2**), with only three genes in common. *ATPase phospholipid transporting 8B3* (*Atp8b3*) was up-regulated in KO rats in both studies, while *Receptor for activated C kinase 1* (*Rack1*) and *SRY-Box Transcription Factor 14* (*Sox14*) were down-regulated in KO rats in both studies.

There were 301 significant DEGs between WT rats of Study 1 and Study 2 (**Table S3**) and 230 significant DEGs between KO rats of Study 1 and Study 2 (**Table S4**), with four genes in common. *Interferon induced protein with tetratricopeptide repeats 3* (*Ifit3*) and *Arginine and serine rich protein 1* (*Rsrp1*) were both up-regulated in Study 2 rats, regardless of genotype; *Kelch domain containing 7A* (*Klhdc7a*) and *Tribbles homolog 2* (*Trib2*) were both down-regulated.

As our focus here was on differences between the studies, we focused IPA analyses on differences between rats of the two studies separately for WT and KO rats. Between WT rats of Study 1 and Study 2, the top difference was an up-regulation in Study 2 of the cholesterol biosynthesis superpathway (Z = 2.24, p = 2e-5; **Supplementary Figure 1a**). Notably, *sterol regulatory element binding factors 1* and *2* (*Srebf1, Srebf2*) were predicted to be activated in Study 2 rats (**Figure 8**). In addition, *glucagon like peptide 2 receptor* (*Glp2r*) was significantly up-regulated in Study 2 rats (log2 = 1.58, p = 0.039; **Table S3**).

**Figure 8.**
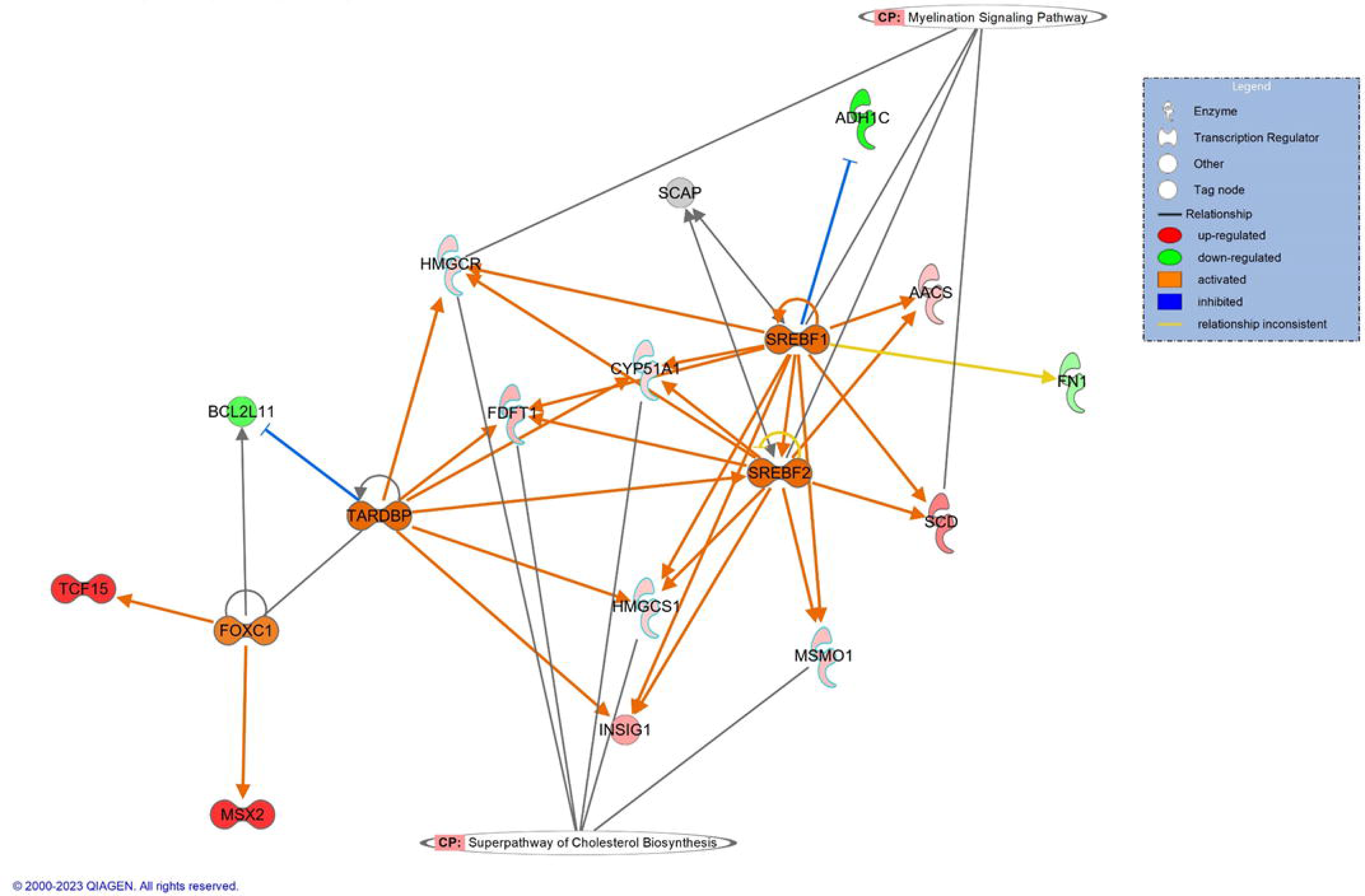
Canonical pathways and upstream regulators between Study 1 and Study 2 in wild-type (WT) rats support changes in environmental stress between the two studies. The top canonical pathway in WT rats was the superpathway for cholesterol biosynthesis, although some genes had connection to the myelination signaling pathway. Notably, there is predicted activation of *sterol regulatory element-binding factors 1* and *2* (*SREBF1, SREBF2*), which have sterol sensing functions. Red and green indicate up- and down-regulated genes in the dataset, respectively. Orange and blue indicate IPA predictions of activity, where orange indicates predicted activation while blue indicates predicted inhibition. Correspondingly, orange arrows suggest activation while blue arrows suggest inhibition; yellow arrows indicate that the prediction is inconsistent with the state of the downstream molecule.

Between KO rats of the studies, the top finding was an up-regulation of the myelination signaling pathway (Z = 1.81, p = 3.59e-7; **Supplementary Figure 1b**). Analysis examining the effect of genotype for each study suggests possible differences in inflammatory response (**Supplementary Figure 2**), but the evidence suggests that the hypothalamus is not the correct tissue to understand genotype-driven differences.

## Discussion

In the current study, there were no differences in the adiposity measures between WT and KO females, contradicting our previous results (13). Upon closer inspection of the data, we determined that in Study 2 WT rats consumed a greater amount of food, which led to an increased body weight and fat mass—WT rats had increased in size without a corresponding change in KO rats. We hypothesize that this difference is due to a significant change in the environment between the two studies, where the environment in Study 1 was likely more stressful. Our CORT data support this hypothesis, showing that WT rats had increased basal CORT relative to KO rats in Study 1, with no differences in Study 2. Changes in expression of CORT metabolism genes and the hypothalamic RNA-seq data further support the argument for environmental differences between the studies. These findings suggest a role of *Krtcap3* in the stress response. Importantly, this work highlights the need to more broadly consider environmental factors in animal research and how those factors may impact replication and translation to human research.

### A change in environment and circulating CORT levels may have altered the adiposity phenotype

As previously described, *Krtcap3* was first identified as a potential adiposity gene through GWAS analyses (10–12, 18) and we confirmed *in vivo* that decreased expression of *Krtcap3* led to increased adiposity (13). We sought here to repeat the *in vivo* study with additional metabolic phenotyping and tissue collection in order to better understand the function of *Krtcap3* in adiposity, but failed to replicate our results. There were significant changes in the adiposity measures of WT rats in the current study relative to the previous study, but no changes in *Krtcap3* expression or the nearby obesity-related gene *Adcy3*. Further, the data imply that Study 2 WT rats were proportionally larger, instead of merely fatter, suggesting that Study 1 WT rats had possibly failed to grow appropriately. This prompted us to wonder what caused such a shift between the two studies. We determined that there were two large environmental changes that we had not considered during study design: the COVID-19 pandemic and associated facility shutdowns plus the completion of a nearby construction project. These changes meant that the environment for the rats in Study 2 was much quieter than it had been in Study 1. Environmental noise is known to impact phenotype in rodent studies (19–21), leading us to hypothesize that the change in environment may have affected the adiposity phenotype by impacting the stress levels of the rats.

We first examined serum CORT in the rats and identified several interesting findings. First, in both studies KO rats had lower serum CORT relative to WT rats, but the difference was highly significant only in Study 1. This supported our hypothesis that stress levels may be related to the adiposity phenotype—differences in basal CORT between WT and KO correlated with differences in adiposity. It was initially surprising that a decrease in CORT was associated with an increase in body weight, as most research has correlated high CORT with obesity and poor metabolic health (22, 23). However, rodents are known to exhibit stress-induced hypophagia depending on factors such as sex, stress type and intensity, and age at stress exposure (24–33).

Furthermore, sub-strains of the WKY, the background strain of the *Krtcap3-*KO rats, differ by body weight and plasma CORT levels in the inverse manner. Specifically, Wistar-Kyoto More Immobile (WMI) females have lower body weight and higher basal CORT compared to Wistar-Kyoto Less Immobile females (34, 35). Given this, we suggest that Study 1 rats were smaller due to increased stress and hypophagia while KO rats were unaffected, presenting an adiposity phenotype in Study 1 that did not manifest when neither genotype was exposed to a stressful environment in Study 2. We propose that the genotype-driven difference we previously saw in adiposity was rather a genotype-driven difference in environmental stress response that had an indirect impact on body weight and fat mass.

This difference in stress response is supported by the distinct CORT response at euthanasia between WT and KO rats: when euthanized second, WT rats have a minimal CORT increase, but KO rats experience a spike in their CORT. We initially presumed this was evidence of chronic stress altering CORT secretion in response to acute stress in the Study 1 WT rats (28, 32, 33), but replicating this pattern in Study 2 rats indicated the effect was not related to environment. We propose that *Krtcap3-*KO rats exhibit low basal CORT and are relatively unaffected by low-grade chronic environmental stress relative to the stress-responsive WKY rat (36) but exhibit a heightened response to acute social stress relative to WT rats. Studies in rodents have previously shown strain-specific differences in CORT response following restraint stress plus differences in glucocorticoid receptor (GR) expression (37–39), supporting a role of genetics in basal CORT levels and stress response. Few causal genes, however, have been found. The current work suggests that *Krtcap3* may be a novel stress gene.

### Changes in CORT processing further indicate a change in environment between Study 1 and Study 2

We also considered if there were changes in gene expression that would further support our hypothesis of an environmental change between the studies. We examined expression of genes related to GC function and processing: *Nr3c1, Hsd11*β*1,* and *Hsd11*β*2. Nr3c1* encodes the GR while the hydroxysteroid dehydrogenases isoforms respectively catalyze the activation or de-activation of CORT (40). Specifically, HSD11β1 is a bidirectional low-affinity enzyme that predominantly reactivates 11-dehydrocorticosterone to corticosterone (41), while HSD11β2 is a high affinity enzyme that exclusively oxidizes the biologically active corticosterone to inactive cortisone (42). Alterations of GC metabolism via these genes has been linked to obesity (43–47). We found that in liver tissue, the expression of the genes *Nr3c1, Hsd11*β*1,* and *Hsd11*β*2* was higher in Study 1 than Study 2, with no genotype-driven differences. Since increased GCs have been shown to up-regulate expression of both *Hsd11*β*1* and *Hsd11*β*2* (48), these findings are in agreement with increased prolonged environmental stress-induced levels of CORT in Study 1. In RetroFat, the most striking difference was the higher expression of *Hsd11*β*2* in Study 2 relative to Study 1, with again no differences by genotype, suggesting a decreased availability of the local bioactive glucocorticoids in Study 2. Whether the increased *Nr3c1* expression in both liver and fat tissue in Study 1 compared to Study 2 is in response to the increased stress environment in Study 1 cannot be determined from this study. However, the lack of genotype difference in these findings suggest the suspected effects of *Krtcap3* on stress regulation are not related to GC processing in the liver or adipose.

### Tissue expression supports a possible role of Krtcap3 in stress response

By examining *Krtcap3* expression in multiple tissues between male and female WKY rats, we have identified several tissues of interest for future investigations into the stress hypothesis and for evaluation of the sex differences we had previously identified (13). *Krtcap3* was highly expressed in the pituitary gland and GI tract of both male and female rats, but only had high expression in the male gonads. The high expression of *Krtcap3* in the pituitary gland supports the possibility of a direct role in the HPA axis and stress response. *Krtcap3* is also highly expressed in the gastrointestinal tract of the rats, which supports our previously suggested role along the gut-brain axis (13). The gut-brain axis has repeatedly been linked to behavior in preclinical and clinical models, germ-free status can negatively impact basal and stimulated HPA activity (49), and stress can disrupt intestinal barrier integrity (50, 51). These findings offer promising new leads for *Krtcap3’s* tissue of action, but additional work is still needed.

### RNA-seq results further suggest the effect of environmental stress on WT rats

Despite low expression of *Krtcap3* in the hypothalamus of WKY/NCrl rats, we chose to perform RNA-seq analysis on hypothalamus tissue from rats from Study 1 and Study 2. There were two reasons for this choice: we had limited tissues that had been collected between both studies and of those tissues the hypothalamus is connected to satiety and obesity (16) and to the stress response (17).

Overall, the results from RNA-seq analysis also support our hypothesis that there was a significant difference in the environments between Study 1 and Study 2, but do not shed much light on the pathways *Krtcap3* acts in. In WT rats, IPA analysis demonstrated up-regulation in cholesterol biosynthesis in Study 2 WT. Cholesterol is vital for neuronal signaling, as it is involved in pre- and post-synaptic neurotransmitter signaling and serves as insulation for myelin sheaths (52). It is of real interest that the major hubs of the pathway are involved in positive regulation of cholesterol biosynthesis and storage, specifically as *Srebf1* and *Srebf2* have sterol sensing functions. An additional interest is that there was a major expression difference in *Glp2r* between WT rats of the two studies, as *Glp2r* has been implicated in the regulation of food intake (53).

There was an up-regulation of myelination signaling pathways in Study 2 compared to Study 1 KOs, and reduced myelin thickness has been well-connected to chronic stress (54, 55). Overall the RNA-seq data demonstrates an increase in neural health in Study 2 relative to Study 1. The data also suggest that *Krtcap3* is likely not acting at the level of the hypothalamus. Differences in basal and stress CORT between WT and KO rats, coupled with the high levels of *Krtcap3* in the pituitary gland suggest *Krtcap3* may be acting at the level of the pituitary. Future studies are necessary for confirmation.

## Conclusion

While we sought to replicate the previous adiposity phenotype between WT and *Krtcap3*-KO rats and identify the mechanism by which *Krtcap3* is influencing adiposity, this study instead demonstrates a possible connection between *Krtcap3* and stress response. In the environment of the second study, WT and KO rats had comparable levels of adiposity due to an increase in eating, body weight, and fat mass in the WT rats relative to the first study, without any corresponding changes in the KO rats. Analyses of CORT indicate that KO rats were able to maintain low CORT despite the potentially stressful environment of Study 1, unlike WT rats, and that decreased *Krtcap3* expression may impart resistance to low-grade chronic environmental stress. This may also account for the lack of change in eating behavior in the KO rats between studies. Expression of GR and the hydroxysteroid dehydrogenases in the liver and visceral fat also support significant differences in CORT processing between the two studies, but do not explain the differences we see between WT and KO rats. The results from RNA-seq analysis also indicate a change in environment between the two studies that primarily had an impact on the neuronal health of WT rats. That *Krtcap3* is highly expressed in both the pituitary and the gastrointestinal tract further support a potential role of *Krtcap3* in stress. In total, these data suggest that decreased expression of *Krtcap3* may protect against low-grade, chronic environmental stress and prevent alterations to eating behavior. WT and KO rats also respond to the euthanasia protocol differently, indicating that KO rats may have a larger response to acute psychosocial stressors than WT rats.

Despite these intriguing findings, there are limitations to this study. We cannot entirely eliminate the possibility that the differences we see are noise. Ours would not be the first lab to see phenotypic differences emerge under different environmental conditions or at a different time (56, 57), and many *in vivo* studies have been marked by a reproducibility crisis (58–60). But we argue that seeing a different phenotype under different environmental conditions does not mean either phenotype is false or that the variation is random. Rather, gene by environment interactions influencing phenotype should be expected, as organisms without such phenotypic plasticity would not survive. Failing to consider environmental factors that do not fall directly under experimental design (19, 61–63) and rigorously standardizing animal experiments (64, 65) fails to consider this plasticity.

An additional limitation is that we currently have not identified where *Krtcap3* is acting in the body to influence CORT, nor are we able to draw conclusions on its mechanistic role in the HPA axis. Finally, although previous work demonstrated sex differences on adiposity (13), the current study was conducted only in female rats and we are unable to make any claims on the role of *Krtcap3* in the stress response of male rats.

While we have identified a potential connection between *Krtcap3* and stress that may play a role in obesity, additional studies are required to confirm this connection. Studies are currently underway to clarify the relationship between *Krtcap3* expression and stress response and to confirm if it has downstream consequences for obesity. Future studies will also identify the tissue of action for *Krtcap3*, elucidate the mechanism of *Krtcap3* in the stress response pathway, identify potential protein interaction partners, and investigate the role of *Krtcap3* in stress in male rats.

### Broader impacts

These data point to an important role of *Krtcap3* in the stress response pathway. Serum CORT measurements indicate that decreased *Krtcap3* expression is associated with low basal CORT and resistance to low-grade chronic environmental stress, but an enhanced response to acute social behavioral stressors. High *Krtcap3* expression in the pituitary and gastrointestinal tract further reflects potential roles in the stress response pathway either by the HPA axis, the gut-brain axis, or both.

The possible role of *Krtcap3* in the stress response is an important consideration for the translational aspect of this research, as the environment for human patients is much more complex than the laboratory environment and plays a significant role in obesity development and treatment. Genes for complex diseases such as obesity must be considered within the environment that human patients exist in. Better consideration of the genetics of stress response (66–69), will improve understanding of stress-related diseases, such as obesity (4, 70–73). Although the underlying molecular mechanisms are still unclear, genetic variation in GC sensitivity in patients has been shown to alter obesity progression and sequelae when patients were exposed to stress (74). Obesity results from a complex web of genetic and environmental factors that are difficult to untangle, and it is necessary to understand how adiposity-related genes function in a translationally relevant environment. Realizing that the role of *Krtcap3* in growth and adiposity may be dependent on its function in the stress response opens up a new avenue of research for this gene in particular, but also broadens our understanding of how to investigate the genetics of obesity moving forward.

In addition to our specific findings for *Krtcap3,* an important takeaway from the current study is what it reveals about animal research. Many animal studies only describe the factors that investigators are able to control: the type of housing, the day/night cycles, the type of food, etc. Factors that are not commonly described in published research—but have been shown to have an impact—include who was handling the animals (63), the noise levels within and outside of the facility (19, 20, 61, 75, 76), the light intensity (61), etc. Failing to fully consider the environment can lead investigators to inadvertently misinterpret results and hinder scientific progress (77) and contributes to science’s reproducibility crisis. Moving forward, genetic research using animal models ought to address these issues, as it is only when we consider the phenotype within the broader context of gene-environment interactions that we can fully understand the role of that gene. For example, without this *in vivo* study, we would not have made the connection between *Krtcap3* and stress. How many other genes might have been ignored because of phenotypic plasticity?

### Data Deposition

RNA-seq data will be deposited in GEO

### Author Contributions

AMS designed the study, conducted experiments, ran statistical analysis, analyzed results, created figures, and wrote the manuscript. LSW designed study, oversaw experimental work, and edited manuscript. GG, EG, and OS assisted with experiments. MG, JK, and AMG created the rat knock-out model. EER consulted on stress hypothesis. All authors approved final version of the manuscript.

## Acknowledgements

The authors thank Dr. Aaron Deal for assisting with feed and weigh, Bailey McDonald for assisting with IPGTT experiments, Katherine Przybyl for conducting initial analyses of corticosterone in the rats, Mackenzie Fitzpatrick for helping dissect inbred rats, and Dr. Laura Cox for her guidance on RNA-seq analysis. The authors also acknowledge BioRender for production of the graphical abstract and Azenta for running RNA-seq analysis and analyzing differentially expressed genes. As always, the authors thank the MCW Genotyping Core for their assistance in genotyping the rats.

**Appendix Figure 1.**
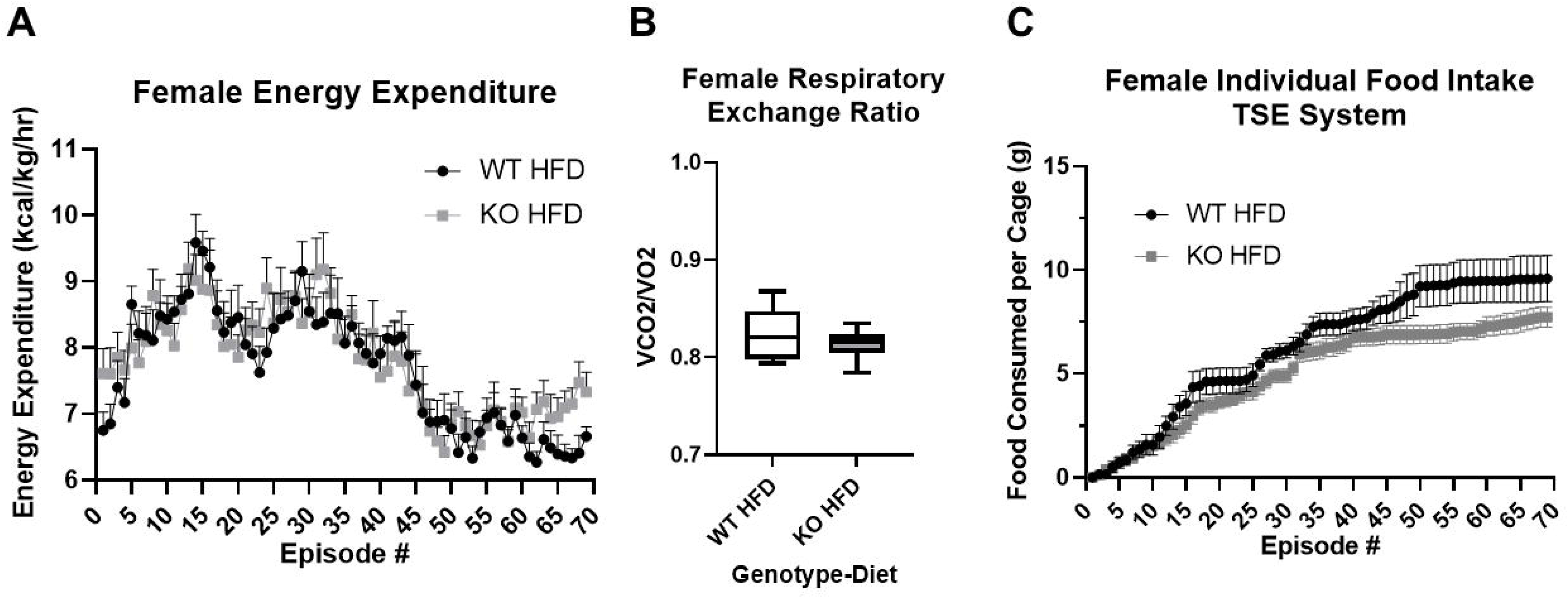
Metabolic rate of Study 2 wild-type (WT) and *Krtcap3* knock-out (KO) rats. Energy expenditure, respiratory exchange ratio, and food consumption were measured over 24 h in TSE Metabolic phenotyping chambers. (A) Energy expenditure. Normalized to individual body weight, there were no differences in energy expenditure between WT (black circle) and KO (gray square) rats. (B) Respiratory exchange ratio. There were no differences in respiratory exchange ratio, a measure of fuel utilization. Finally, (B) Food consumption. There were also no significant differences in food consumption while in the TSE chambers.

**Appendix Figure 2.**
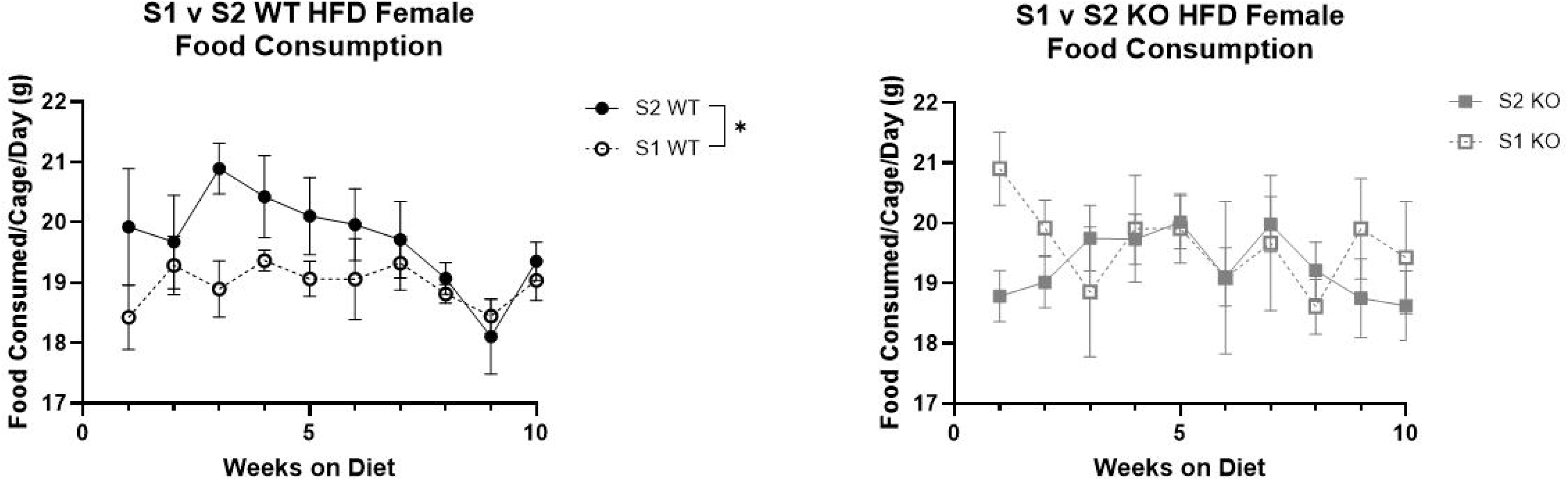
Differences in food consumption between Study 1 (S1) and Study 2 (S2) in wild-type (WT) and *Krtcap3* knock-out (KO) rats during the first 10 weeks on diet. Food intake by cage (two rats) was measured weekly for both studies, and grams of food consumed per cage per day could be calculated. Study 2 WT (black filled circle, solid line) consumed more food than Study 1 WT (black open circle, dashed line) for most of the study. There were no consistent differences in food consumption between Study 2 KO (gray filled circle, solid line) and Study 1 KO (gray empty circle, dashed line). *p < 0.05 represents the effect of study respective to genotype.

**Appendix Figure 3.**
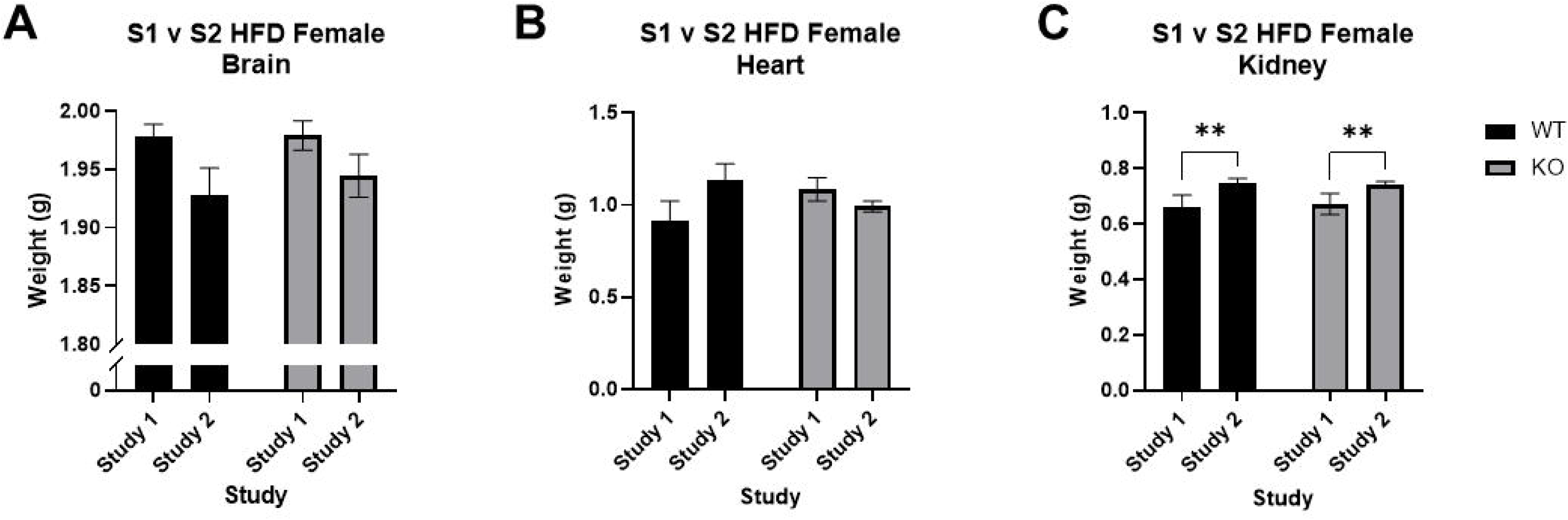
Comparing non-adipose organ weights between Study 1 (S1) and Study 2 (S2) in wild-type (WT) and *Krtcap3* knock-out (KO) rats. Liver could not be included because its full weight was not recorded in Study 1. (A) Brain weights showed no statistically significant differences by genotype or study. (B) Heart weights also showed no significant differences. (C) Kidney weight showed a significant increase in Study 2 animals, regardless of genotype. *p < 0.05 represents an effect of study respective to genotype, **p < 0.01 represents a main effect of study.

**Appendix Figure 4.**
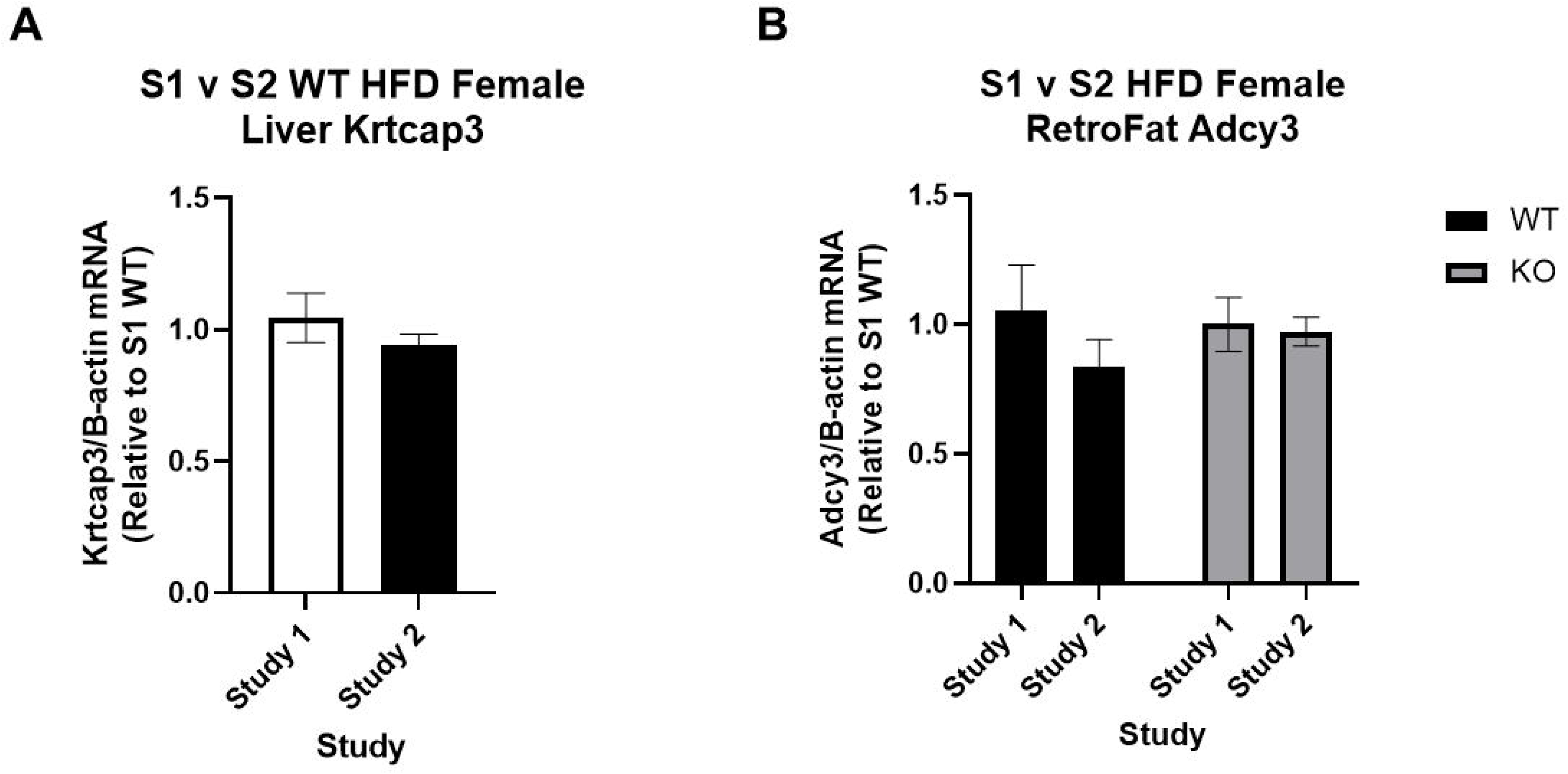
No changes in expression of (A) *Krtcap3* or (B) *Adcy3* between Study 1 (S1) and Study 2 (S2). (A) There were no differences in liver *Krtcap3* expression between Study 1 and Study 2 wild-type (WT) rats. (B) There were no changes in *Adcy3* expression by study in either WT or *Krtcap3* knock-out (KO) rats. Further, there were no differences in expression between WT and KO rats.

**Appendix Table 1.**
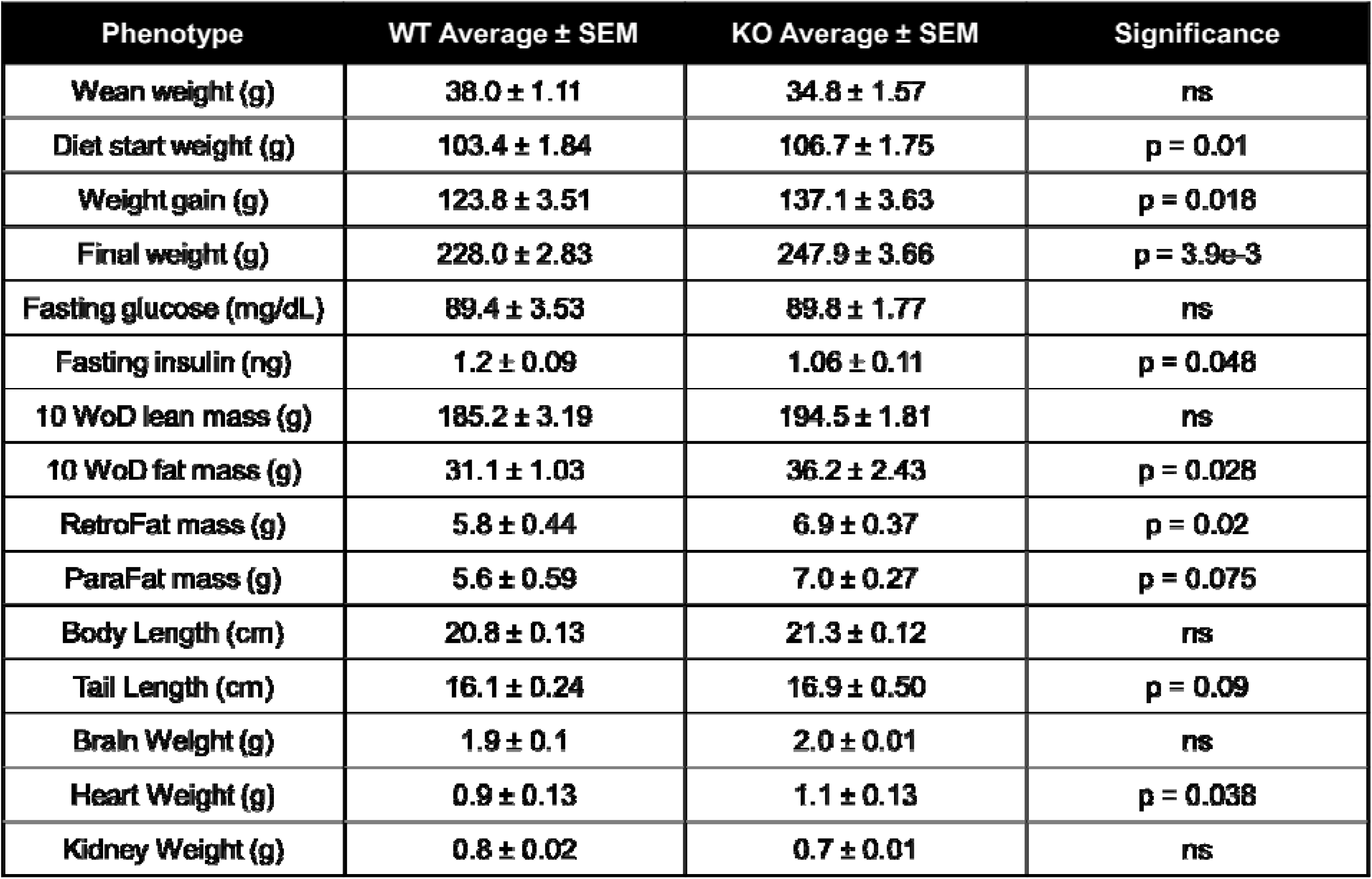
Body size, adiposity, and non-adipose organ weights between wild-type (WT) and *Krtcap3* knock-out (KO) female rats from Study 1. Weeks on diet (WoD), area under the curve (AUC), retroperitoneal fat (RetroFat), and parametrial fat (ParaFat). Body length was measured from nose to anus, and tail length was measured from anus to end of tail. Values are given as the mean average ± standard error. Significance is indicated in the final column.

