## Supplementary Table 1 for "Changes in Environmental Stress over COVID-19 Pandemic Likely Contributed to Failure to Replicate Adiposity Phenotype Associated with *Krtcap3*"

| ID | Symbol | Log2 Fold Change | p-value |
| --- | --- | --- | --- |
| ENSRNOG00000010633 | ACSL1 | -0.16506 | 0.046366 |
| ENSRNOG00000019180 | ACSL4 | 0.295502 | 0.003985 |
| ENSRNOG00000002232 | AFF1 | -0.38301 | 0.042985 |
| ENSRNOG00000016696 | ANGPT2 | 0.702321 | 0.013354 |
| ENSRNOG00000019534 | AP2A2 | 0.164268 | 0.045341 |
| ENSRNOG00000000166 | APEX2 | -0.41719 | 0.010223 |
| ENSRNOG00000014521 | ARMC8 | 0.124786 | 0.049313 |
| ENSRNOG00000045649 | ARRDC3 | -0.25272 | 0.01494 |
| ENSRNOG00000024975 | ATP8B3 | 0.401379 | 0.029334 |
| ENSRNOG00000001971 | BBX | -0.31312 | 0.031766 |
| ENSRNOG00000017459 | C1QL3 | 0.429205 | 0.008971 |
| ENSRNOG00000005669 | CA8 | 0.438849 | 0.042101 |
| ENSRNOG00000009954 | CACUL1 | 0.178775 | 0.045386 |
| ENSRNOG00000021781 | CAMK1 | 0.234859 | 0.023387 |
| ENSRNOG00000058938 | CAMKV | 0.323247 | 0.020695 |
| ENSRNOG00000047453 | CASD1 | 0.226892 | 0.000873 |
| ENSRNOG00000025643 | CCDC13 | -0.28657 | 0.03991 |
| ENSRNOG00000003893 | Ccnjl | -0.45429 | 0.034267 |
| ENSRNOG00000002141 | CD200 | 0.235647 | 0.025629 |
| ENSRNOG00000006000 | CDK12 | -0.19898 | 0.044858 |
| ENSRNOG00000025012 | CHAT | -1.09553 | 0.025277 |
| ENSRNOG00000009974 | COQ3 | 0.257441 | 0.029051 |
| ENSRNOG00000015397 | CPNE7 | 0.368675 | 0.038984 |
| ENSRNOG00000012443 | CPT2 | -0.32661 | 0.014413 |
| ENSRNOG00000005330 | CREBBP | -0.19948 | 0.034105 |
| ENSRNOG00000037146 | CTXN2 | 0.278534 | 0.049866 |
| ENSRNOG00000015980 | DCLRE1C | -0.40979 | 0.020468 |
| ENSRNOG00000057078 | DDIT4 | -0.29501 | 0.007096 |
| ENSRNOG00000022368 | DDX49 | -0.1958 | 0.043357 |
| ENSRNOG00000001106 | DENR | 0.177232 | 0.033529 |
| ENSRNOG00000008062 | DQX1 | -0.46248 | 0.008616 |
| ENSRNOG00000050190 | ENG | -0.25415 | 0.025011 |
| ENSRNOG00000061147 | ENTREP1 | -0.48492 | 0.041995 |
| ENSRNOG00000018336 | EPS8L2 | -0.52309 | 0.019474 |
| ENSRNOG00000014801 | EXOG | 0.21885 | 0.044824 |
| ENSRNOG00000021067 | FXYD7 | 0.395678 | 0.014909 |
| ENSRNOG00000004645 | GALNT5 | -2.12692 | 0.049673 |
| ENSRNOG00000004360 | GEMIN2 | -0.3334 | 0.017731 |
| ENSRNOG00000048980 | GNG2 | 0.204662 | 0.037772 |
| ENSRNOG00000021020 | GPHA2 | -2.71382 | 0.036939 |
| ENSRNOG00000010731 | GPM6A | 0.207494 | 0.009114 |
| ENSRNOG00000060137 | GPR173 | 0.234864 | 0.036295 |
| ENSRNOG00000018991 | GSN | -0.24141 | 0.041872 |
| ENSRNOG00000049952 | GUCD1 | -0.36113 | 0.017225 |
| ENSRNOG00000059362 | HAS3 | -0.58531 | 0.019347 |
| ENSRNOG00000023969 | HERC6 | -0.35509 | 0.023253 |
| ENSRNOG00000018796 | HERPUD1 | -0.14011 | 0.039701 |
| ENSRNOG00000023299 | HFM1 | 0.42398 | 0.033632 |
| ENSRNOG00000001686 | HLCS | -0.25307 | 0.029993 |
| ENSRNOG00000020601 | HMG20B | -0.26386 | 0.017053 |
| ENSRNOG00000001338 | HPD | -0.98568 | 0.011743 |
| ENSRNOG00000006227 | IFIH1 | -0.43591 | 0.023133 |
| ENSRNOG00000027430 | IKZF2 | -0.66409 | 0.005993 |
| ENSRNOG00000012151 | ITPRIPL1 | -0.55395 | 0.039426 |
| ENSRNOG00000050343 | Jmy | -0.26702 | 0.010905 |
| ENSRNOG00000013312 | KCNT2 | 0.454761 | 0.047473 |
| ENSRNOG00000015669 | KCTD11 | -0.47546 | 0.0385 |
| ENSRNOG00000011846 | KRT28 | -1.31354 | 0.019549 |
| ENSRNOG00000031579 | LOC100363469 | 0.864143 | 0.042378 |
| ENSRNOG00000033015 | LOC100910150 | -0.70861 | 0.038132 |
| ENSRNOG00000048686 | LOC103690742 (includes others) | 0.26827 | 0.003617 |
| ENSRNOG00000055292 | LOC120102832 | -0.5732 | 0.019337 |
| ENSRNOG00000031669 | LPP | -0.83306 | 0.01682 |
| ENSRNOG00000001727 | LSG1 | -0.25035 | 0.038783 |
| ENSRNOG00000003268 | MAML1 | -0.29013 | 0.036668 |
| ENSRNOG00000014021 | MATN4 | -0.52941 | 0.018955 |
| ENSRNOG00000021702 | MCMDC2 | 0.242981 | 0.046424 |
| ENSRNOG00000013674 | MEGF10 | -0.24123 | 0.018235 |
| ENSRNOG00000045574 | MORN3 | 0.602703 | 0.021484 |
| ENSRNOG00000020991 | MS4A6A | 0.708903 | 0.038596 |
| ENSRNOG00000029042 | MT-ND6 | 0.280495 | 0.043948 |
| ENSRNOG00000000075 | MTF2 | -0.20683 | 0.044841 |
| ENSRNOG00000061928 | MYO1E | -0.28618 | 0.036876 |
| ENSRNOG00000013911 | NAGK | 0.189602 | 0.046593 |
| ENSRNOG00000013562 | NDFIP1 | 0.129371 | 0.026745 |
| ENSRNOG00000040040 | NDUFAF6 | 0.322023 | 0.017727 |
| ENSRNOG00000004505 | NFIC | -0.2979 | 0.04191 |
| ENSRNOG00000023258 | NFKB1 | -0.14997 | 0.045767 |
| ENSRNOG00000007390 | NFKBIA | -0.37043 | 0.043162 |
| ENSRNOG00000005392 | NGFR | -0.58281 | 0.015025 |
| ENSRNOG00000018095 | NKIRAS2 | -0.39945 | 0.015165 |
| ENSRNOG00000016156 | NPTXR | 0.362407 | 0.005357 |
| ENSRNOG00000007607 | NR4A1 | -0.49041 | 0.043202 |
| ENSRNOG00000010392 | NRG1 | -0.35161 | 0.0207 |
| ENSRNOG00000017037 | OTUD3 | -0.31898 | 0.02433 |
| ENSRNOG00000029756 | P2RY13 | 0.410482 | 0.00738 |
| ENSRNOG00000007574 | PADI2 | -0.19917 | 0.019584 |
| ENSRNOG00000011526 | PCSK6 | -0.43789 | 0.003442 |
| ENSRNOG00000002517 | PDC | 1.081436 | 0.006685 |
| ENSRNOG00000008748 | PEX2 | -0.23274 | 0.026174 |
| ENSRNOG00000020371 | Pgap2 | -0.21517 | 0.025595 |
| ENSRNOG00000010873 | PITHD1 | 0.112086 | 0.028013 |
| ENSRNOG00000012095 | PKIA | 0.189728 | 0.004981 |
| ENSRNOG00000000811 | PKIB | 0.560976 | 0.033309 |
| ENSRNOG00000026662 | PLEKHF2 | 0.226387 | 0.047723 |
| ENSRNOG00000047917 | PLSCR5 | 0.824419 | 0.008542 |
| ENSRNOG00000020906 | POLA2 | -0.37307 | 0.036516 |
| ENSRNOG00000004719 | PP2D1 | 0.264511 | 0.049436 |
| ENSRNOG00000028733 | PRKAR1B | 0.140475 | 0.019227 |
| ENSRNOG00000004288 | PTRH2 | 0.422199 | 0.000353 |
| ENSRNOG00000014705 | RBM20 | -1.06577 | 0.021956 |
| ENSRNOG00000050413 | SAMD15 | -0.40971 | 0.039053 |
| ENSRNOG00000002104 | SCAF4 | -0.20014 | 0.031251 |
| ENSRNOG00000005018 | SCN2A | 0.182236 | 0.042493 |
| ENSRNOG00000001337 | Setd1a | -0.23846 | 0.025803 |
| ENSRNOG00000028198 | Sh2b3 | -0.22589 | 0.045302 |
| ENSRNOG00000027468 | SLC6A15 | 0.13373 | 0.032242 |
| ENSRNOG00000023404 | SLC9B2 | 0.385889 | 0.044163 |
| ENSRNOG00000049689 | SMIM18 | 0.375615 | 0.028977 |
| ENSRNOG00000008829 | SORBS3 | -0.19145 | 0.041872 |
| ENSRNOG00000022084 | SOX14 | -1.17672 | 0.026841 |
| ENSRNOG00000008209 | ST3GAL1 | -0.32713 | 0.038641 |
| ENSRNOG00000001960 | Sult1d1 | 0.483942 | 0.000532 |
| ENSRNOG00000016133 | SUMO1 | 0.136673 | 0.045684 |
| ENSRNOG00000008157 | SYN2 | 0.213486 | 0.044858 |
| ENSRNOG00000050547 | SYNGR2 | -0.21161 | 0.034094 |
| ENSRNOG00000011823 | TFAP2B | -1.62792 | 0.019922 |
| ENSRNOG00000050500 | TOB2 | -0.3302 | 0.003573 |
| ENSRNOG00000004100 | TRIB1 | -0.72464 | 0.033065 |
| ENSRNOG00000014981 | USP14 | 0.115315 | 0.024753 |
| ENSRNOG00000048606 | Vma21 | 0.371029 | 0.039733 |
| ENSRNOG00000014564 | YIPF5 | 0.156737 | 0.021834 |
| ENSRNOG00000014785 | YKT6 | 0.167136 | 0.039284 |
| ENSRNOG00000004588 | ZBED4 | -0.36775 | 0.015895 |
| ENSRNOG00000010436 | ZBTB17 | -0.18755 | 0.048392 |
| ENSRNOG00000014380 | Zfp788 | 0.203273 | 0.019708 |
| ENSRNOG00000011627 | ZFR | 0.108189 | 0.025065 |
| ENSRNOG00000027436 | ZNF324 | -0.32962 | 0.027429 |
| ENSRNOG00000019802 | ZNF428 | 0.198043 | 0.04369 |
| ENSRNOG00000047526 | ZNF526 | -0.22158 | 0.042892 |
| ENSRNOG00000009990 | Zranb2 | 0.14734 | 0.04837 |
| ENSRNOG00000018160 | ZSWIM5 | -0.25421 | 0.044259 |

**Table S4.** Significantly differentially expressed genes (DEGs) between wild-type (WT) and *Krtcap3* knock-out (KO) rats of Study 1.
