## Supplementary Table 2 for "Changes in Environmental Stress over COVID-19 Pandemic Likely Contributed to Failure to Replicate Adiposity Phenotype Associated with *Krtcap3*"

| ID | Symbol | Log2 Fold Change | p-value |
| --- | --- | --- | --- |
| ENSRNOG00000046871 | 2310039H08Rik | -0.47514 | 0.020224 |
| ENSRNOG00000008862 | ABCG4 | 0.19054 | 0.021496 |
| ENSRNOG00000037080 | ADAMTS17 | 0.712468 | 0.016879 |
| ENSRNOG00000010840 | ADAMTSL3 | -0.54102 | 0.017258 |
| ENSRNOG00000002922 | ADORA2B | -0.42686 | 0.002977 |
| ENSRNOG00000021256 | ADRA1D | 1.012932 | 0.031818 |
| ENSRNOG00000013720 | AEBP1 | -0.79777 | 0.026377 |
| ENSRNOG00000045738 | AK4 | -0.2547 | 0.025055 |
| ENSRNOG00000017672 | Akr1c14 | -0.93502 | 0.001106 |
| ENSRNOG00000047023 | ALDH1L1 | -0.40966 | 0.037177 |
| ENSRNOG00000014610 | ANPEP | -0.53125 | 0.031878 |
| ENSRNOG00000017469 | ANXA1 | -0.61121 | 0.038485 |
| ENSRNOG00000003829 | AP3S1 | 0.172283 | 0.029126 |
| ENSRNOG00000043059 | APLF | -0.3056 | 0.023113 |
| ENSRNOG00000011648 | AQP1 | -1.27373 | 0.005586 |
| ENSRNOG00000016043 | AQP4 | -0.27385 | 0.035357 |
| ENSRNOG00000025502 | ARHGEF37 | 0.28438 | 0.046319 |
| ENSRNOG00000026060 | ARSI | -1.0508 | 0.023292 |
| ENSRNOG00000046858 | Arxes1/Arxes2 | -0.21698 | 0.028145 |
| ENSRNOG00000049714 | ASAP3 | -0.58256 | 0.034137 |
| ENSRNOG00000009197 | ASB4 | -0.35733 | 0.043745 |
| ENSRNOG00000000415 | ASF1A | 0.178151 | 0.042273 |
| ENSRNOG00000017913 | ATG16L1 | 0.158549 | 0.032011 |
| ENSRNOG00000000885 | Auts2l1 | 0.334071 | 0.028241 |
| ENSRNOG00000004049 | BAIAP2 | -0.41491 | 0.023645 |
| ENSRNOG00000055962 | BGN | -0.48964 | 0.009975 |
| ENSRNOG00000021745 | BHLHE22 | -0.70448 | 0.011501 |
| ENSRNOG00000002322 | C1orf115 | 0.388276 | 0.019428 |
| ENSRNOG00000011718 | C1RL | -0.84271 | 0.020196 |
| ENSRNOG00000011971 | C1S | -0.3341 | 0.018062 |
| ENSRNOG00000026600 | CAPS2 | -0.71437 | 0.030739 |
| ENSRNOG00000031129 | CARM1 | 0.192398 | 0.006278 |
| ENSRNOG00000025895 | CAVIN2 | -0.40482 | 0.000484 |
| ENSRNOG00000028581 | CCDC138 | -0.40753 | 0.009981 |
| ENSRNOG00000002052 | CCDC80 | -0.34115 | 0.02592 |
| ENSRNOG00000048848 | Ccdc9 | 0.142304 | 0.031881 |
| ENSRNOG00000008697 | CCN3 | -0.64333 | 3.48E-05 |
| ENSRNOG00000057058 | CD300A | 0.778043 | 0.010837 |
| ENSRNOG00000006094 | CD44 | -0.88824 | 0.002475 |
| ENSRNOG00000059500 | Cdkn1c | -0.73873 | 0.033195 |
| ENSRNOG00000003908 | Cep128 | -0.28495 | 0.037861 |
| ENSRNOG00000053230 | CEP20 | -0.18446 | 0.048557 |
| ENSRNOG00000043410 | CFAP300 | -0.43408 | 0.016628 |
| ENSRNOG00000029911 | CILP | -0.94342 | 0.021432 |
| ENSRNOG00000020622 | CILP2 | 1.054156 | 0.031105 |
| ENSRNOG00000046607 | CITED4 | 0.739499 | 0.014773 |
| ENSRNOG00000054495 | CLDN2 | -3.29114 | 0.000325 |
| ENSRNOG00000026870 | CLIC6 | -0.64278 | 0.002649 |
| ENSRNOG00000011260 | CMBL | -0.23665 | 0.037582 |
| ENSRNOG00000022838 | CNKSR1 | -0.61464 | 0.038261 |
| ENSRNOG00000047307 | CNTFR | 0.204227 | 0.017127 |
| ENSRNOG00000001229 | COL18A1 | -0.2618 | 0.032174 |
| ENSRNOG00000003349 | COL23A1 | -1.0321 | 0.003618 |
| ENSRNOG00000003357 | COL3A1 | -0.73244 | 0.013692 |
| ENSRNOG00000015365 | COL4A3 | 0.622832 | 0.02199 |
| ENSRNOG00000014851 | Col4a4 | 0.613305 | 0.026118 |
| ENSRNOG00000003736 | COL5A2 | -0.29025 | 0.000919 |
| ENSRNOG00000001254 | COL6A2 | -0.53757 | 0.01886 |
| ENSRNOG00000019648 | COL6A3 | -0.6158 | 0.018813 |
| ENSRNOG00000010841 | COL8A2 | -0.95081 | 0.006094 |
| ENSRNOG00000016366 | COLEC12 | -0.57535 | 0.011712 |
| ENSRNOG00000021155 | CTSK | -0.59447 | 0.04769 |
| ENSRNOG00000022256 | CXCL10 | -0.95693 | 0.044524 |
| ENSRNOG00000017511 | DCDC2 | -0.60778 | 0.009243 |
| ENSRNOG00000000842 | DDAH2 | -0.28875 | 0.01113 |
| ENSRNOG00000026748 | DENND2A | -0.25535 | 0.037527 |
| ENSRNOG00000030157 | DHX16 | 0.176972 | 0.033563 |
| ENSRNOG00000038916 | DRAM1 | -0.63409 | 0.043342 |
| ENSRNOG00000032307 | DSEL | -0.30711 | 0.013578 |
| ENSRNOG00000036798 | DUSP3 | 0.173505 | 0.036816 |
| ENSRNOG00000015596 | EFHD1 | 0.194913 | 0.0399 |
| ENSRNOG00000020588 | EFNA4 | -0.82239 | 0.038666 |
| ENSRNOG00000005123 | EMC2 | -0.14678 | 0.03635 |
| ENSRNOG00000008246 | EMILIN1 | -0.2841 | 0.023025 |
| ENSRNOG00000018310 | ENO4 | 0.279702 | 0.032099 |
| ENSRNOG00000045952 | EPHB4 | -0.3425 | 0.041367 |
| ENSRNOG00000048187 | EPOP | 0.585325 | 0.027407 |
| ENSRNOG00000057855 | F5 | -2.2538 | 0.00023 |
| ENSRNOG00000049075 | FABP5 | -0.25003 | 0.007746 |
| ENSRNOG00000023008 | FAM131C | 0.242973 | 0.045604 |
| ENSRNOG00000011316 | FAM167A | -0.68653 | 0.04394 |
| ENSRNOG00000011750 | FAM180A | -0.95634 | 0.036724 |
| ENSRNOG00000001198 | FAM222A | 0.469067 | 0.004733 |
| ENSRNOG00000043219 | FBN2 | 0.656313 | 0.048725 |
| ENSRNOG00000054515 | FGD6 | 0.37643 | 0.045002 |
| ENSRNOG00000015941 | FKBP10 | -0.26345 | 0.039428 |
| ENSRNOG00000009886 | FKBP14 | -0.34886 | 0.013907 |
| ENSRNOG00000005478 | FKBP9 | -0.29356 | 0.025573 |
| ENSRNOG00000024917 | FLACC1 | -0.63495 | 0.02121 |
| ENSRNOG00000020421 | FNDC7 | -0.42558 | 0.020236 |
| ENSRNOG00000047446 | FOXC2 | -1.22776 | 0.008668 |
| ENSRNOG00000016189 | FRMD5 | 0.170769 | 0.033094 |
| ENSRNOG00000007765 | FRZB | -0.75988 | 0.000417 |
| ENSRNOG00000033100 | FTH1 | -0.495 | 0.016649 |
| ENSRNOG00000003470 | FUNDC1 | -0.18614 | 0.03923 |
| ENSRNOG00000060599 | GABRB3 | -0.20563 | 0.013131 |
| ENSRNOG00000010183 | GASK1B | -0.37407 | 0.027571 |
| ENSRNOG00000027040 | GGACT | -0.34249 | 0.039377 |
| ENSRNOG00000042152 | GIPC2 | -0.76062 | 0.02489 |
| ENSRNOG00000027837 | Gm14569 | -0.74914 | 0.047573 |
| ENSRNOG00000013339 | GMEB2 | 0.33215 | 0.023074 |
| ENSRNOG00000018969 | GPATCH4 | 0.330939 | 0.007645 |
| ENSRNOG00000002413 | GPC4 | -0.47532 | 0.042686 |
| ENSRNOG00000056457 | GPD1 | 0.216682 | 0.021771 |
| ENSRNOG00000013171 | GRM2 | 0.535579 | 0.013085 |
| ENSRNOG00000038891 | GRXCR1 | -1.0991 | 0.009544 |
| ENSRNOG00000013484 | Gsta1 | -0.38832 | 0.00603 |
| ENSRNOG00000030449 | Gsta4 | -0.21684 | 0.023004 |
| ENSRNOG00000054017 | H4C11 | -0.44248 | 0.006605 |
| ENSRNOG00000016967 | HFE | -0.63851 | 0.032034 |
| ENSRNOG00000007027 | HGF | -1.08665 | 0.019586 |
| ENSRNOG00000000451 | HLA-DQA1 | -1.23447 | 0.007326 |
| ENSRNOG00000032708 | HLA-DQB1 | -0.9942 | 0.0327 |
| ENSRNOG00000032844 | HLA-DRA | -1.02817 | 0.029705 |
| ENSRNOG00000016122 | HMGCR | -0.19447 | 0.027416 |
| ENSRNOG00000014887 | HOMEZ | -0.40511 | 0.016211 |
| ENSRNOG00000014964 | HP | -0.72552 | 0.008017 |
| ENSRNOG00000017659 | HS3ST2 | -0.53259 | 0.013155 |
| ENSRNOG00000020533 | HTRA1 | 0.227161 | 0.031394 |
| ENSRNOG00000026605 | Ifi27l2b | -1.04589 | 0.001514 |
| ENSRNOG00000021966 | IL17RD | 0.493688 | 0.010898 |
| ENSRNOG00000048924 | ISLR | -0.98307 | 0.015528 |
| ENSRNOG00000007905 | ITGA7 | 0.235985 | 0.045881 |
| ENSRNOG00000048449 | ITGB3 | -0.75237 | 0.034628 |
| ENSRNOG00000016023 | KANK1 | 0.185031 | 0.016888 |
| ENSRNOG00000010643 | KANK2 | -0.26012 | 0.039666 |
| ENSRNOG00000004918 | KCNA4 | 0.397863 | 0.048663 |
| ENSRNOG00000009872 | KCNH2 | 0.242129 | 0.046119 |
| ENSRNOG00000018018 | KCNIP2 | -1.00808 | 0.038232 |
| ENSRNOG00000033796 | KCNJ5 | -0.45869 | 0.007008 |
| ENSRNOG00000012494 | KCTD14 | -0.43855 | 0.048609 |
| ENSRNOG00000046829 | KDR | -0.22925 | 0.042768 |
| ENSRNOG00000013661 | KIF26A | 0.367414 | 0.026015 |
| ENSRNOG00000014029 | KLHL13 | -0.20995 | 0.015079 |
| ENSRNOG00000025273 | L3mbtl4 | -0.78015 | 0.043145 |
| ENSRNOG00000010552 | LAMTOR3 | 0.186571 | 0.044101 |
| ENSRNOG00000006865 | LAPTM4A | -0.14714 | 0.006804 |
| ENSRNOG00000014532 | LBP | -1.29585 | 0.012997 |
| ENSRNOG00000024470 | LMAN1 | -0.17382 | 0.046012 |
| ENSRNOG00000019551 | LOC100360846/Psmb6 | -0.13594 | 0.041885 |
| ENSRNOG00000031579 | LOC100363469 | 0.902046 | 0.049165 |
| ENSRNOG00000010891 | LRRCC1 | -0.28523 | 0.043222 |
| ENSRNOG00000051408 | MAGT1 | -0.20433 | 0.032482 |
| ENSRNOG00000015101 | MAK | -0.63526 | 0.029517 |
| ENSRNOG00000002848 | MAOA | -0.14098 | 0.023149 |
| ENSRNOG00000018781 | MAP1S | 0.129362 | 0.04377 |
| ENSRNOG00000003952 | MAP3K19 | -0.57629 | 0.040239 |
| ENSRNOG00000007271 | MAP3K9 | 0.337069 | 0.011167 |
| ENSRNOG00000004583 | MB | 1.394987 | 0.035863 |
| ENSRNOG00000008610 | MBIP | -0.30307 | 0.015518 |
| ENSRNOG00000020029 | MCRIP2 | 0.330952 | 0.036851 |
| ENSRNOG00000039107 | MFRP | -1.12885 | 0.003324 |
| ENSRNOG00000024954 | MGAT5B | 0.398339 | 0.036933 |
| ENSRNOG00000000175 | MIER2 | 0.198148 | 0.037424 |
| ENSRNOG00000007924 | MIOS | -0.20691 | 0.033234 |
| ENSRNOG00000035471 | mir-124 | 1.194762 | 0.031857 |
| ENSRNOG00000005382 | Mroh4 | 3.960398 | 0.006068 |
| ENSRNOG00000010540 | MRPL45 | 0.212788 | 0.018575 |
| ENSRNOG00000059381 | MRPS6 | -0.20756 | 0.049956 |
| ENSRNOG00000018355 | MSX2 | -1.13603 | 0.017125 |
| ENSRNOG00000013641 | MYO7A | 0.399564 | 0.035984 |
| ENSRNOG00000011754 | MYOM2 | -0.57416 | 0.010334 |
| ENSRNOG00000054506 | NCCRP1 | 1.033337 | 0.010718 |
| ENSRNOG00000018577 | NEIL1 | -0.36337 | 0.004699 |
| ENSRNOG00000016245 | NETO2 | 0.273276 | 0.035069 |
| ENSRNOG00000026055 | NEUROD6 | -0.95616 | 0.014022 |
| ENSRNOG00000049083 | NICN1 | 0.120957 | 0.047739 |
| ENSRNOG00000002461 | NID1 | -0.30618 | 0.029971 |
| ENSRNOG00000013085 | NKAIN3 | -0.74356 | 0.022849 |
| ENSRNOG00000013408 | NPAS2 | 0.511211 | 0.024039 |
| ENSRNOG00000019184 | NPR3 | -0.67479 | 0.03936 |
| ENSRNOG00000003741 | NPTX1 | -0.24548 | 0.043732 |
| ENSRNOG00000001585 | NRIP1 | 0.260582 | 0.022976 |
| ENSRNOG00000046053 | NUDT11 | -0.26845 | 0.047149 |
| ENSRNOG00000009862 | OLFM1 | 0.295539 | 0.041775 |
| ENSRNOG00000019883 | PAK4 | -0.26385 | 0.027766 |
| ENSRNOG00000054212 | PDE1A | -0.35977 | 0.047665 |
| ENSRNOG00000008543 | PDLIM2 | -0.89604 | 0.016334 |
| ENSRNOG00000012658 | PDLIM3 | -0.57062 | 0.049246 |
| ENSRNOG00000015406 | PGM5 | -0.34008 | 0.032957 |
| ENSRNOG00000014668 | PHAF1 | 0.187844 | 0.021026 |
| ENSRNOG00000004019 | PHLDA1 | -0.35025 | 0.001011 |
| ENSRNOG00000002171 | PHLDB2 | -0.31882 | 0.042575 |
| ENSRNOG00000000145 | PIK3R3 | -0.20775 | 0.047813 |
| ENSRNOG00000048676 | PIP5KL1 | 0.24631 | 0.043313 |
| ENSRNOG00000004205 | PKDCC | -0.38139 | 0.019466 |
| ENSRNOG00000008747 | PLEKHA5 | 0.239378 | 0.009435 |
| ENSRNOG00000016479 | PLEKHG4 | -0.57162 | 0.043648 |
| ENSRNOG00000023085 | PMEL | 0.292448 | 0.042197 |
| ENSRNOG00000012664 | POLR1E | 0.380241 | 0.014156 |
| ENSRNOG00000017416 | PPIC | -0.69715 | 0.049561 |
| ENSRNOG00000046535 | PPM1M | -0.41844 | 0.032673 |
| ENSRNOG00000025350 | PPP1R13L | 0.581064 | 0.002496 |
| ENSRNOG00000003120 | PRELP | -0.57432 | 0.010521 |
| ENSRNOG00000048723 | PROS1 | -0.31507 | 0.035825 |
| ENSRNOG00000019041 | PSME1 | -0.34751 | 0.002584 |
| ENSRNOG00000027432 | R3HDML | 1.024698 | 0.019126 |
| ENSRNOG00000020067 | RAB40C | 0.289322 | 0.019486 |
| ENSRNOG00000028872 | RAI14 | -0.41415 | 0.012619 |
| ENSRNOG00000010219 | RALGDS | 0.137326 | 0.019587 |
| ENSRNOG00000056151 | RASGRF2 | -0.4354 | 0.010357 |
| ENSRNOG00000010595 | RBM45 | 0.204074 | 0.037254 |
| ENSRNOG00000002074 | REST | -0.27227 | 0.027918 |
| ENSRNOG00000017604 | RIPOR1 | 0.133548 | 0.044598 |
| ENSRNOG00000060162 | Rnu6-909 | -1.16546 | 0.009812 |
| ENSRNOG00000037247 | RRAS | -0.43138 | 0.024521 |
| ENSRNOG00000037687 | RSPO2 | -0.63222 | 0.031534 |
| ENSRNOG00000000636 | RTKN2 | 0.521529 | 0.016885 |
| ENSRNOG00000004222 | S100G | 1.844482 | 0.015581 |
| ENSRNOG00000026116 | SALL3 | -0.28415 | 0.042557 |
| ENSRNOG00000020342 | SAMD11 | -0.62463 | 0.016931 |
| ENSRNOG00000013552 | SCD | -0.38853 | 0.010788 |
| ENSRNOG00000031203 | SCFD1 | -0.2213 | 0.015182 |
| ENSRNOG00000010617 | SCUBE1 | -0.38294 | 0.038267 |
| ENSRNOG00000006631 | SEMA3E | 0.465089 | 0.043723 |
| ENSRNOG00000014650 | SEMA4G | 0.22978 | 0.0465 |
| ENSRNOG00000049244 | SEPSECS | -0.20857 | 0.036654 |
| ENSRNOG00000009465 | SFRP2 | 0.59148 | 0.048408 |
| ENSRNOG00000003038 | SFT2D2 | -0.3276 | 0.033711 |
| ENSRNOG00000028238 | Sh3bgr | -0.6685 | 0.00943 |
| ENSRNOG00000054400 | SH3BP5L | 0.157315 | 0.033758 |
| ENSRNOG00000005522 | SH3YL1 | -0.37617 | 0.016368 |
| ENSRNOG00000011503 | SHB | 0.373545 | 0.020736 |
| ENSRNOG00000059057 | SHLD2 | -0.29807 | 0.044906 |
| ENSRNOG00000019791 | SIPA1L2 | 0.211655 | 0.047291 |
| ENSRNOG00000019118 | SLC13A3 | -0.42587 | 0.039342 |
| ENSRNOG00000021644 | SLC15A3 | 0.496381 | 0.043685 |
| ENSRNOG00000002832 | SLC16A2 | -0.59665 | 0.001898 |
| ENSRNOG00000018215 | SLC22A6 | -1.16938 | 0.006672 |
| ENSRNOG00000038001 | Slc25a1 | -0.17252 | 0.028993 |
| ENSRNOG00000004351 | SLC25A29 | -0.23783 | 0.024178 |
| ENSRNOG00000016827 | SLC38A3 | 0.173613 | 0.019196 |
| ENSRNOG00000059463 | SLC39A1 | -0.14707 | 0.032879 |
| ENSRNOG00000045524 | SLC39A3 | 0.273064 | 0.031119 |
| ENSRNOG00000011723 | SLC44A3 | -0.78218 | 0.022321 |
| ENSRNOG00000018822 | SLC5A5 | -0.7545 | 0.003234 |
| ENSRNOG00000005126 | SLC66A3 | -0.3718 | 0.044044 |
| ENSRNOG00000010210 | SLC7A11 | -0.50674 | 0.00294 |
| ENSRNOG00000015567 | SLC9A2 | -0.56839 | 0.032704 |
| ENSRNOG00000032798 | SLCO3A1 | 0.19046 | 0.046443 |
| ENSRNOG00000009296 | SNAPC1 | 0.24956 | 0.015293 |
| ENSRNOG00000023548 | SNED1 | -0.31198 | 0.042391 |
| ENSRNOG00000061361 | Snord46 | 0.996723 | 0.027912 |
| ENSRNOG00000022084 | SOX14 | -1.20506 | 0.006601 |
| ENSRNOG00000054086 | SP5 | -1.46082 | 0.00453 |
| ENSRNOG00000029862 | SPC24 | -0.96733 | 0.042085 |
| ENSRNOG00000057881 | SPPL2B | 0.15212 | 0.033883 |
| ENSRNOG00000006733 | SRGAP2 | 0.166303 | 0.028069 |
| ENSRNOG00000038948 | STAB2 | 0.659005 | 0.034981 |
| ENSRNOG00000017051 | STIMATE-MUSTN1 | 0.186757 | 0.034693 |
| ENSRNOG00000019294 | STK16 | 0.196883 | 0.017916 |
| ENSRNOG00000014030 | SYNM | -0.45238 | 0.000306 |
| ENSRNOG00000017628 | TAGLN | -0.46953 | 0.039997 |
| ENSRNOG00000052477 | TBRG4 | 0.16026 | 0.041097 |
| ENSRNOG00000049020 | TCEAL3 | -0.15742 | 0.04733 |
| ENSRNOG00000011387 | TET3 | 0.243571 | 0.021698 |
| ENSRNOG00000002414 | TFCP2L1 | -0.53359 | 0.042276 |
| ENSRNOG00000045829 | THBS1 | -0.67719 | 0.001082 |
| ENSRNOG00000053753 | Thsd4 | -0.79778 | 0.001845 |
| ENSRNOG00000006941 | THUMPD3 | -0.15666 | 0.047036 |
| ENSRNOG00000057290 | TMEM168 | -0.3377 | 0.03448 |
| ENSRNOG00000047783 | TMEM200A | -0.2903 | 0.039842 |
| ENSRNOG00000028085 | TMEM255A | -0.41243 | 0.041577 |
| ENSRNOG00000023340 | TMEM72 | -1.11751 | 0.008141 |
| ENSRNOG00000018943 | TNNC1 | 0.598161 | 0.031208 |
| ENSRNOG00000033734 | TNNT2 | -1.14488 | 0.006282 |
| ENSRNOG00000024849 | TOR1AIP2 | -0.18796 | 0.048375 |
| ENSRNOG00000012609 | TRDN | -1.17096 | 0.025852 |
| ENSRNOG00000016465 | TRMU | -0.21195 | 0.049692 |
| ENSRNOG00000014714 | TRPV6 | -0.23498 | 0.030807 |
| ENSRNOG00000003786 | TSPAN6 | -0.25809 | 0.035167 |
| ENSRNOG00000013877 | TTC23 | -0.61178 | 0.04619 |
| ENSRNOG00000023895 | UBE2U | -0.66675 | 0.042441 |
| ENSRNOG00000000692 | UNG | -0.55772 | 0.032854 |
| ENSRNOG00000003785 | USP43 | 1.073818 | 0.03755 |
| ENSRNOG00000028774 | VGLL3 | -1.22737 | 0.002068 |
| ENSRNOG00000018087 | VIM | -0.51424 | 0.006655 |
| ENSRNOG00000012001 | VPS37A | 0.229141 | 0.012712 |
| ENSRNOG00000011918 | VSX2 | -0.98489 | 0.037099 |
| ENSRNOG00000010031 | VTN | -0.24251 | 0.048548 |
| ENSRNOG00000021628 | WDR89 | -0.48982 | 0.036312 |
| ENSRNOG00000006217 | WIZ | 0.200018 | 0.019759 |
| ENSRNOG00000005781 | WNT16 | -0.62111 | 0.036003 |
| ENSRNOG00000008168 | WNT5B | -0.67985 | 0.016171 |
| ENSRNOG00000000918 | ZBED5 | 0.344241 | 0.005424 |
| ENSRNOG00000056716 | ZBTB20 | -0.28389 | 0.024557 |
| ENSRNOG00000027115 | ZC2HC1C | -0.36124 | 0.043956 |
| ENSRNOG00000025634 | ZFHX2 | 0.274286 | 0.03519 |
| ENSRNOG00000019477 | ZMYND15 | -0.77764 | 0.022891 |
| ENSRNOG00000013379 | ZNF22 | -0.20651 | 0.021817 |

**Table S5.** Significantly differentially expressed genes (DEGs) between wild-type (WT) and *Krtcap3* knock-out (KO) rats of Study 2.
