## Supplementary Table 3 for "Changes in Environmental Stress over COVID-19 Pandemic Likely Contributed to Failure to Replicate Adiposity Phenotype Associated with *Krtcap3*"

| ID | Symbol | Log2 Fold Change | p-value |
| --- | --- | --- | --- |
| ENSRNOG00000000967 | AACS | 0.248141 | 0.03116 |
| ENSRNOG00000008424 | AAGAB | -0.2462 | 0.016245 |
| ENSRNOG00000015666 | ABCC12 | 1.212016 | 0.007014 |
| ENSRNOG00000020433 | ACTN4 | -0.10633 | 0.032012 |
| ENSRNOG00000003206 | ACTR3 | 0.123455 | 0.040781 |
| ENSRNOG00000000563 | ADAMTS14 | -0.75904 | 0.039018 |
| ENSRNOG00000009892 | ADAMTS15 | -0.83831 | 0.03436 |
| ENSRNOG00000037080 | ADAMTS17 | -0.57653 | 0.023706 |
| ENSRNOG00000061484 | ADAMTS2 | 0.714686 | 0.006595 |
| ENSRNOG00000012436 | ADH1C | -1.17483 | 0.043467 |
| ENSRNOG00000021256 | ADRA1D | -1.36536 | 0.048602 |
| ENSRNOG00000052572 | Aff2 | 0.185747 | 0.028097 |
| ENSRNOG00000059445 | AIFM2 | 0.241648 | 0.015602 |
| ENSRNOG00000012841 | ALG11 | 0.15825 | 0.038026 |
| ENSRNOG00000000166 | APEX2 | -0.50351 | 0.022902 |
| ENSRNOG00000036880 | ARL5C | 1.323919 | 0.017817 |
| ENSRNOG00000018693 | ASGR1 | 0.684557 | 0.049722 |
| ENSRNOG00000017913 | ATG16L1 | -0.16937 | 0.047894 |
| ENSRNOG00000013209 | BARHL1 | 1.262197 | 0.021344 |
| ENSRNOG00000002117 | BARHL2 | 0.838142 | 0.025372 |
| ENSRNOG00000016551 | BCL2L11 | -0.74103 | 0.035219 |
| ENSRNOG00000021745 | BHLHE22 | 1.244086 | 0.000334 |
| ENSRNOG00000021773 | BOP1 | -0.18766 | 0.039543 |
| ENSRNOG00000055564 | C11orf96 | -0.33711 | 0.045008 |
| ENSRNOG00000002963 | C1QL1 | 0.663314 | 0.009684 |
| ENSRNOG00000001621 | C2CD2 | -0.34962 | 0.017745 |
| ENSRNOG00000002191 | C4orf19 | -0.61079 | 0.015808 |
| ENSRNOG00000001289 | C7orf50 | 0.240478 | 0.033256 |
| ENSRNOG00000002916 | CA4 | -0.62216 | 0.040659 |
| ENSRNOG00000033531 | CACNA2D1 | 0.311797 | 0.007765 |
| ENSRNOG00000002572 | CACYBP | 0.141994 | 0.025496 |
| ENSRNOG00000016977 | CALB2 | 0.628752 | 0.019473 |
| ENSRNOG00000020239 | CAPN15 | -0.2199 | 0.042054 |
| ENSRNOG00000025518 | CARMIL3 | 0.337528 | 0.039725 |
| ENSRNOG00000047453 | CASD1 | 0.178108 | 0.030045 |
| ENSRNOG00000002905 | CCDC103 | 1.14313 | 0.035857 |
| ENSRNOG00000022953 | Ccdc163 | -0.39125 | 0.040351 |
| ENSRNOG00000042711 | CCDC188 | -1.0771 | 0.025288 |
| ENSRNOG00000001930 | Ccdc50 | 0.208993 | 0.03637 |
| ENSRNOG00000004057 | CCDC88A | 0.159922 | 0.030337 |
| ENSRNOG00000048848 | Ccdc9 | -0.18913 | 0.004721 |
| ENSRNOG00000003893 | Ccnjl | -0.56567 | 0.008769 |
| ENSRNOG00000002141 | CD200 | 0.253988 | 0.024439 |
| ENSRNOG00000045558 | CD34 | -0.37565 | 0.0341 |
| ENSRNOG00000024799 | CD93 | -0.59514 | 0.000835 |
| ENSRNOG00000004743 | CDCP1 | -1.78363 | 0.048526 |
| ENSRNOG00000033087 | CDH23 | 0.587957 | 0.015237 |
| ENSRNOG00000006000 | CDK12 | -0.2512 | 0.017279 |
| ENSRNOG00000022899 | CDK15 | -1.59127 | 0.010371 |
| ENSRNOG00000045771 | CHL1 | 0.27133 | 0.00052 |
| ENSRNOG00000046000 | Clasrp | 0.202551 | 0.012322 |
| ENSRNOG00000007014 | CNKSR2 | 0.294218 | 0.00589 |
| ENSRNOG00000027175 | CNPY4 | -0.26048 | 0.046582 |
| ENSRNOG00000029556 | Cntnap5c | 0.345412 | 0.004188 |
| ENSRNOG00000019918 | COASY | 0.23153 | 0.048367 |
| ENSRNOG00000000463 | COL11A2 | 0.205039 | 0.004076 |
| ENSRNOG00000015365 | COL4A3 | -0.62579 | 0.036757 |
| ENSRNOG00000014851 | Col4a4 | -0.68942 | 0.019095 |
| ENSRNOG00000009974 | COQ3 | 0.314587 | 0.020046 |
| ENSRNOG00000012826 | CREB3L2 | -0.27324 | 0.036692 |
| ENSRNOG00000006622 | CRY1 | -0.30238 | 0.027601 |
| ENSRNOG00000048025 | CSTF2 | 0.16588 | 0.046637 |
| ENSRNOG00000030790 | CTNND1 | -0.22001 | 0.027144 |
| ENSRNOG00000004257 | CTPS2 | 0.17662 | 0.047404 |
| ENSRNOG00000013589 | CXCL12 | -0.23048 | 0.040409 |
| ENSRNOG00000000196 | CYP19A1 | -1.03698 | 0.033432 |
| ENSRNOG00000007234 | CYP51A1 | 0.174663 | 0.034602 |
| ENSRNOG00000033099 | DCC | 0.345986 | 0.012588 |
| ENSRNOG00000012771 | DDX41 | -0.34678 | 0.041909 |
| ENSRNOG00000011716 | DEGS2 | -0.42412 | 0.031697 |
| ENSRNOG00000046050 | DENND1C | -0.45301 | 0.009344 |
| ENSRNOG00000026748 | DENND2A | 0.187852 | 0.043586 |
| ENSRNOG00000010794 | DENND3 | -0.22127 | 0.043424 |
| ENSRNOG00000030157 | DHX16 | -0.15527 | 0.045308 |
| ENSRNOG00000007054 | DHX33 | -0.25714 | 0.046693 |
| ENSRNOG00000014011 | DLL4 | -0.60173 | 0.013487 |
| ENSRNOG00000050742 | DNMBP | -0.26529 | 0.039231 |
| ENSRNOG00000010311 | DZIP1 | 0.20799 | 0.04322 |
| ENSRNOG00000029333 | ECHDC2 | 0.444618 | 0.032328 |
| ENSRNOG00000010815 | ELF2 | -0.19005 | 0.049507 |
| ENSRNOG00000019162 | EMC9 | 0.242153 | 0.035209 |
| ENSRNOG00000050190 | ENG | -0.25503 | 0.036677 |
| ENSRNOG00000043006 | EPM2AIP1 | 0.135341 | 0.029123 |
| ENSRNOG00000018336 | EPS8L2 | -0.80144 | 0.019424 |
| ENSRNOG00000058186 | ERRFI1 | -0.20715 | 0.020703 |
| ENSRNOG00000014125 | EVI2B | 0.658171 | 0.012188 |
| ENSRNOG00000014476 | Evl | 0.167907 | 0.019591 |
| ENSRNOG00000010396 | EYA3 | -0.18708 | 0.036957 |
| ENSRNOG00000011800 | F3 | -0.51886 | 0.001561 |
| ENSRNOG00000001198 | FAM222A | -0.66277 | 0.028089 |
| ENSRNOG00000018327 | FANK1 | 0.687055 | 0.045216 |
| ENSRNOG00000033912 | Fcho1 | 0.196274 | 0.022092 |
| ENSRNOG00000021314 | FDFT1 | 0.308304 | 0.000534 |
| ENSRNOG00000003578 | FEM1C | -0.42214 | 0.047138 |
| ENSRNOG00000010213 | FGD5 | 0.267647 | 0.039885 |
| ENSRNOG00000017225 | FHIP2A | -0.24441 | 0.012846 |
| ENSRNOG00000004699 | FIBIN | -0.45127 | 0.009734 |
| ENSRNOG00000000940 | Flt1 | -0.23653 | 0.032897 |
| ENSRNOG00000003510 | FMO2 | -0.60295 | 0.020256 |
| ENSRNOG00000014288 | FN1 | -0.42244 | 0.005332 |
| ENSRNOG00000009284 | FOXA1 | 1.568344 | 0.025937 |
| ENSRNOG00000029089 | FOXR1 | 1.17581 | 0.045289 |
| ENSRNOG00000039152 | FREM3 | -1.04237 | 0.033107 |
| ENSRNOG00000033100 | FTH1 | 0.454469 | 0.048991 |
| ENSRNOG00000026964 | GAS8 | 0.265598 | 0.046762 |
| ENSRNOG00000018091 | GDI2 | 0.105868 | 0.029874 |
| ENSRNOG00000015860 | GIPR | 0.612225 | 0.045687 |
| ENSRNOG00000003683 | GLP2R | 1.576542 | 0.038735 |
| ENSRNOG00000019838 | GMFG | 0.620083 | 0.029337 |
| ENSRNOG00000015051 | GOLGA7B | 0.255801 | 0.031407 |
| ENSRNOG00000021020 | GPHA2 | -2.24187 | 0.048595 |
| ENSRNOG00000017702 | GPLD1 | 0.262885 | 0.026808 |
| ENSRNOG00000009540 | GPR3 | -0.90635 | 0.038507 |
| ENSRNOG00000024636 | GPR85 | 0.296331 | 0.013321 |
| ENSRNOG00000014615 | GRK6 | 0.166182 | 0.017452 |
| ENSRNOG00000027722 | H1-10 | 0.380451 | 0.04358 |
| ENSRNOG00000047321 | HBA1/HBA2 | 0.744493 | 0.025432 |
| ENSRNOG00000023299 | HFM1 | 0.551283 | 0.037958 |
| ENSRNOG00000016122 | HMGCR | 0.205202 | 0.02332 |
| ENSRNOG00000016552 | HMGCS1 | 0.208046 | 0.011502 |
| ENSRNOG00000017659 | HS3ST2 | 0.586354 | 0.014797 |
| ENSRNOG00000030877 | HTR2C | 0.336876 | 0.033615 |
| ENSRNOG00000022839 | IFIT3 | 0.67184 | 0.02902 |
| ENSRNOG00000017277 | IGSF6 | -0.7875 | 0.008333 |
| ENSRNOG00000013480 | ING2 | -0.29703 | 0.032633 |
| ENSRNOG00000006859 | INSIG1 | 0.381645 | 0.009364 |
| ENSRNOG00000012151 | ITPRIPL1 | -0.6231 | 0.032137 |
| ENSRNOG00000016023 | KANK1 | -0.22792 | 0.043341 |
| ENSRNOG00000007160 | KAT14 | -0.36059 | 0.003678 |
| ENSRNOG00000062002 | KCNA3 | 0.387044 | 0.017727 |
| ENSRNOG00000013463 | KCNJ8 | -0.58433 | 0.028864 |
| ENSRNOG00000013781 | KCNQ5 | 0.450114 | 0.007972 |
| ENSRNOG00000050200 | KDM3B | -0.18753 | 0.037492 |
| ENSRNOG00000018141 | KIAA0319 | 0.208141 | 0.032451 |
| ENSRNOG00000013661 | KIF26A | -0.59504 | 0.013064 |
| ENSRNOG00000016299 | KLF4 | -0.54736 | 0.027833 |
| ENSRNOG00000018867 | KLHDC7A | -0.59379 | 0.02689 |
| ENSRNOG00000014029 | KLHL13 | 0.221712 | 0.049847 |
| ENSRNOG00000003756 | LANCL3 | -0.4369 | 0.027739 |
| ENSRNOG00000025448 | LIMD2 | 0.283024 | 0.035473 |
| ENSRNOG00000012988 | LIX1 | 0.161822 | 0.048859 |
| ENSRNOG00000016161 | LMAN2 | -0.17644 | 0.042012 |
| ENSRNOG00000017019 | LMX1B | 1.480016 | 0.03285 |
| ENSRNOG00000000891 | LOC102548389 | 0.202586 | 0.046262 |
| ENSRNOG00000048686 | LOC103690742 (includes others) | 0.26849 | 0.005008 |
| ENSRNOG00000024796 | LRRC47 | 0.141197 | 0.040725 |
| ENSRNOG00000029798 | LRRC4C | 0.196909 | 0.023788 |
| ENSRNOG00000026466 | LRRTM3 | 0.27996 | 0.039954 |
| ENSRNOG00000003453 | LYPD1 | 0.379032 | 0.014012 |
| ENSRNOG00000021400 | MAGEE2 | 0.812783 | 0.030225 |
| ENSRNOG00000014971 | MAS1 | 1.174025 | 0.031593 |
| ENSRNOG00000008610 | MBIP | 0.347938 | 0.00948 |
| ENSRNOG00000000536 | MDGA1 | 0.285676 | 0.04946 |
| ENSRNOG00000016210 | MICAL2 | -0.82471 | 0.01937 |
| ENSRNOG00000003613 | MID1 | 0.352249 | 0.045569 |
| ENSRNOG00000007924 | MIOS | 0.215406 | 0.024665 |
| ENSRNOG00000035529 | mir-135 | 1.259524 | 0.003641 |
| ENSRNOG00000035617 | mir-186 | 0.428921 | 0.025392 |
| ENSRNOG00000035583 | mir-208 | -1.10219 | 0.007967 |
| ENSRNOG00000031093 | MOV10L1 | 1.137123 | 0.041302 |
| ENSRNOG00000005382 | Mroh4 | -4.2695 | 0.00092 |
| ENSRNOG00000032297 | MSMO1 | 0.260466 | 0.011248 |
| ENSRNOG00000018355 | MSX2 | 1.058218 | 0.028812 |
| ENSRNOG00000004685 | MTA3 | 0.125999 | 0.038719 |
| ENSRNOG00000002697 | MTMR1 | -0.21547 | 0.007003 |
| ENSRNOG00000022929 | MTMR12 | -0.43185 | 0.015301 |
| ENSRNOG00000015236 | MYBBP1A | -0.12754 | 0.042312 |
| ENSRNOG00000002215 | MYLK | -0.57043 | 0.04973 |
| ENSRNOG00000032994 | MYOM3 | 1.137781 | 0.013237 |
| ENSRNOG00000021525 | NBEAL1 | 0.331019 | 0.01229 |
| ENSRNOG00000002126 | NCAM2 | 0.224516 | 0.02743 |
| ENSRNOG00000056678 | NCKAP5L | -0.1964 | 0.047473 |
| ENSRNOG00000010620 | NDC1 | -0.25043 | 0.006578 |
| ENSRNOG00000009577 | NDST4 | 0.770434 | 0.003444 |
| ENSRNOG00000028417 | NEUROD2 | 1.252017 | 0.01055 |
| ENSRNOG00000008449 | NEUROD4 | -1.78265 | 0.008864 |
| ENSRNOG00000026055 | NEUROD6 | 1.194233 | 0.014381 |
| ENSRNOG00000006611 | NOSTRIN | -0.4665 | 0.040127 |
| ENSRNOG00000016156 | NPTXR | 0.216012 | 0.046767 |
| ENSRNOG00000007607 | NR4A1 | -0.63655 | 0.028842 |
| ENSRNOG00000001585 | NRIP1 | -0.39441 | 0.0059 |
| ENSRNOG00000051837 | NWD2 | 0.56352 | 0.007016 |
| ENSRNOG00000036668 | OGFOD3 | 0.340184 | 0.031562 |
| ENSRNOG00000024043 | ORC6 | 0.368464 | 0.034241 |
| ENSRNOG00000029756 | P2RY13 | 0.462731 | 0.00834 |
| ENSRNOG00000024729 | PAX5 | 1.50903 | 0.005046 |
| ENSRNOG00000046264 | Pcdhb7 | 0.507506 | 0.004345 |
| ENSRNOG00000011526 | PCSK6 | -0.36426 | 0.018712 |
| ENSRNOG00000002517 | PDC | 1.005345 | 0.048437 |
| ENSRNOG00000002169 | PDCL2 | 0.691015 | 0.041965 |
| ENSRNOG00000014443 | PDE5A | -0.54203 | 0.03836 |
| ENSRNOG00000050932 | PFDN4 | 0.265679 | 0.036615 |
| ENSRNOG00000014668 | PHAF1 | -0.17948 | 0.032466 |
| ENSRNOG00000000121 | PIGV | -0.28069 | 0.026138 |
| ENSRNOG00000008798 | PIPOX | 0.498869 | 0.024923 |
| ENSRNOG00000010873 | PITHD1 | 0.141681 | 0.023958 |
| ENSRNOG00000012095 | PKIA | 0.205257 | 0.018736 |
| ENSRNOG00000000811 | PKIB | 0.592349 | 0.043616 |
| ENSRNOG00000024904 | PLA2G4E | 0.441084 | 0.03589 |
| ENSRNOG00000016479 | PLEKHG4 | 0.815977 | 0.014888 |
| ENSRNOG00000023140 | Pnldc1 | 0.887929 | 0.030104 |
| ENSRNOG00000012664 | POLR1E | -0.33835 | 0.037568 |
| ENSRNOG00000017503 | PPARGC1B | -0.33648 | 0.046294 |
| ENSRNOG00000025350 | PPP1R13L | -0.58729 | 0.003932 |
| ENSRNOG00000050483 | PRADC1 | -0.41446 | 0.011834 |
| ENSRNOG00000008674 | PRDM11 | 0.275595 | 0.039442 |
| ENSRNOG00000014180 | PRICKLE4 | 0.260006 | 0.020162 |
| ENSRNOG00000029865 | PRSS56 | 1.779383 | 0.000932 |
| ENSRNOG00000017730 | PSMD1 | 0.121551 | 0.019495 |
| ENSRNOG00000001210 | Pwp2 | -0.18048 | 0.027068 |
| ENSRNOG00000039754 | RAB7B | 0.43236 | 0.038539 |
| ENSRNOG00000000956 | RASL11A | -1.44066 | 0.020314 |
| ENSRNOG00000015780 | RCN2 | 0.20306 | 0.029029 |
| ENSRNOG00000003283 | RCSD1 | -1.03337 | 0.024016 |
| ENSRNOG00000027919 | RDH13 | -0.21934 | 0.04891 |
| ENSRNOG00000061774 | RGD1560986 | 0.718271 | 0.031313 |
| ENSRNOG00000005515 | RHBDL3 | 0.262931 | 0.041935 |
| ENSRNOG00000054385 | RHEBL1 | -0.43248 | 0.025493 |
| ENSRNOG00000017568 | RIT2 | 0.163428 | 0.047196 |
| ENSRNOG00000009658 | RNF19A | -0.21858 | 0.02246 |
| ENSRNOG00000016387 | Rpl34 (includes others) | -0.15874 | 0.008517 |
| ENSRNOG00000017309 | RSRP1 | 0.133072 | 0.036406 |
| ENSRNOG00000017060 | RYR2 | 0.306732 | 0.017615 |
| ENSRNOG00000019987 | SCAND1 | 0.167328 | 0.041659 |
| ENSRNOG00000013552 | SCD | 0.516875 | 0.015185 |
| ENSRNOG00000020083 | SCLY | -0.25712 | 0.049586 |
| ENSRNOG00000029342 | SCN7A | -0.5486 | 0.022363 |
| ENSRNOG00000001337 | Setd1a | -0.17739 | 0.045559 |
| ENSRNOG00000012536 | SGMS1 | -0.24898 | 0.020994 |
| ENSRNOG00000015967 | SH3BGRL3 | 0.206654 | 0.021122 |
| ENSRNOG00000005522 | SH3YL1 | 0.475486 | 0.007245 |
| ENSRNOG00000019826 | SIL1 | -0.23464 | 0.0329 |
| ENSRNOG00000016123 | SIM1 | -0.29842 | 0.027581 |
| ENSRNOG00000007250 | SIX4 | -0.62456 | 0.029621 |
| ENSRNOG00000019996 | SLC16A1 | -0.24701 | 0.025819 |
| ENSRNOG00000038001 | Slc25a1 | 0.238414 | 0.028725 |
| ENSRNOG00000017075 | SLC35E2B | -0.32818 | 0.049882 |
| ENSRNOG00000004604 | SLC38A10 | -0.23263 | 0.031988 |
| ENSRNOG00000027767 | SLC38A5 | 0.546801 | 0.019483 |
| ENSRNOG00000011981 | SLC39A13 | -0.18449 | 0.048952 |
| ENSRNOG00000002240 | SLC49A4 | 0.161828 | 0.041115 |
| ENSRNOG00000028844 | SLC9A5 | 0.218053 | 0.031307 |
| ENSRNOG00000023404 | SLC9B2 | 0.35109 | 0.046163 |
| ENSRNOG00000007377 | SLIT3 | -0.38473 | 0.048697 |
| ENSRNOG00000039877 | Smim11 | 0.33419 | 0.040154 |
| ENSRNOG00000054374 | Smim29 | 0.216591 | 0.049389 |
| ENSRNOG00000008656 | SNCA | 0.600684 | 0.035741 |
| ENSRNOG00000061361 | Snord46 | -1.05393 | 0.026005 |
| ENSRNOG00000046874 | SNX9 | -0.25124 | 0.049693 |
| ENSRNOG00000058173 | Sox3 | -0.76961 | 0.031183 |
| ENSRNOG00000021032 | SPHK2 | -0.18485 | 0.026896 |
| ENSRNOG00000008209 | ST3GAL1 | -0.42971 | 0.021334 |
| ENSRNOG00000056243 | ST7 | 0.272189 | 0.046967 |
| ENSRNOG00000001090 | STARD13 | -0.28051 | 0.039286 |
| ENSRNOG00000011705 | STMN2 | 0.174885 | 0.040264 |
| ENSRNOG00000015670 | STX7 | 0.173244 | 0.048724 |
| ENSRNOG00000001960 | Sult1d1 | 0.357041 | 0.016092 |
| ENSRNOG00000026315 | TAF7L | 1.090579 | 0.032607 |
| ENSRNOG00000053574 | TARBP1 | -0.27543 | 0.039545 |
| ENSRNOG00000014750 | TASOR | 0.158693 | 0.049146 |
| ENSRNOG00000000012 | TCF15 | 1.48736 | 0.024973 |
| ENSRNOG00000003878 | THSD7B | 0.425844 | 0.007944 |
| ENSRNOG00000020189 | TINF2 | 0.214083 | 0.038976 |
| ENSRNOG00000012349 | TLNRD1 | -0.33341 | 0.036498 |
| ENSRNOG00000000073 | TMED5 | -0.25771 | 0.018991 |
| ENSRNOG00000021517 | TMEM231 | -0.25443 | 0.049316 |
| ENSRNOG00000023340 | TMEM72 | 1.441863 | 0.004593 |
| ENSRNOG00000016731 | Tpm2 | -0.37507 | 0.025812 |
| ENSRNOG00000004110 | Trib2 | -0.27849 | 0.045816 |
| ENSRNOG00000015829 | TRPM1 | 0.900617 | 0.048146 |
| ENSRNOG00000013053 | TRPM6 | -1.2181 | 0.010488 |
| ENSRNOG00000014714 | TRPV6 | 0.263428 | 0.041833 |
| ENSRNOG00000008758 | TSPAN18 | 0.390502 | 0.0343 |
| ENSRNOG00000001219 | Tspear | 0.496343 | 0.039275 |
| ENSRNOG00000024526 | TTC34 | 0.725597 | 0.035077 |
| ENSRNOG00000048411 | UHRF1 | 0.523008 | 0.026023 |
| ENSRNOG00000008237 | UNC13B | 0.267911 | 0.042367 |
| ENSRNOG00000014981 | USP14 | 0.129777 | 0.009162 |
| ENSRNOG00000017714 | USP3 | 0.294704 | 0.033668 |
| ENSRNOG00000026880 | USP38 | -0.2129 | 0.047708 |
| ENSRNOG00000016477 | VANGL1 | -0.6542 | 0.046372 |
| ENSRNOG00000021156 | VEGFB | 0.195158 | 0.049403 |
| ENSRNOG00000018808 | VIP | -1.62365 | 0.038535 |
| ENSRNOG00000025539 | VPS13A | 0.24574 | 0.024378 |
| ENSRNOG00000011918 | VSX2 | 1.245292 | 0.033549 |
| ENSRNOG00000003825 | WDR75 | -0.22584 | 0.032712 |
| ENSRNOG00000012929 | WSB1 | 0.219972 | 0.044651 |
| ENSRNOG00000011634 | XKR6 | 0.557368 | 0.011198 |
| ENSRNOG00000056716 | ZBTB20 | 0.414961 | 0.024325 |
| ENSRNOG00000006943 | ZCCHC10 | 0.42042 | 0.044802 |
| ENSRNOG00000022686 | ZDHHC2 | 0.234576 | 0.025337 |
| ENSRNOG00000049137 | Zfp35 | -0.37883 | 0.011114 |
| ENSRNOG00000053366 | Zfp59 | 0.294234 | 0.012482 |
| ENSRNOG00000023920 | Zfp791 | 0.493891 | 0.034468 |
| ENSRNOG00000004109 | ZFPM2 | 0.384829 | 0.039689 |
| ENSRNOG00000022414 | ZNF142 | -0.18809 | 0.012501 |
| ENSRNOG00000042283 | ZNF267 | 0.290906 | 0.035809 |
| ENSRNOG00000010985 | ZNF410 | -0.17134 | 0.02791 |
| ENSRNOG00000047526 | ZNF526 | -0.23633 | 0.039568 |
| ENSRNOG00000002682 | ZNF692 | 0.160461 | 0.026252 |
| ENSRNOG00000022391 | ZNF711 | 0.322391 | 0.043028 |

**Table S6.** Significantly differentially expressed genes (DEGs) between Study 1 and Study 2 wild-type (WT) rats.
