## Supplementary Table 4 for "Changes in Environmental Stress over COVID-19 Pandemic Likely Contributed to Failure to Replicate Adiposity Phenotype Associated with *Krtcap3*"

| ID | Symbol | Log2 Fold Change | p-value |
| --- | --- | --- | --- |
| ENSRNOG00000004147 | Abca8a | 0.200727 | 0.021286 |
| ENSRNOG00000015002 | ABHD15 | 0.713846 | 0.011128 |
| ENSRNOG00000010840 | ADAMTSL3 | -0.72617 | 0.005121 |
| ENSRNOG00000017672 | Akr1c14 | -0.7827 | 0.039641 |
| ENSRNOG00000016696 | ANGPT2 | -0.79288 | 0.001417 |
| ENSRNOG00000014610 | ANPEP | -0.67499 | 0.016316 |
| ENSRNOG00000022771 | ARHGAP23 | 0.144742 | 0.034801 |
| ENSRNOG00000012563 | ARHGAP29 | -0.32186 | 0.015513 |
| ENSRNOG00000028090 | ARHGEF18 | 0.149699 | 0.022159 |
| ENSRNOG00000007733 | ARHGEF9 | -0.13659 | 0.039605 |
| ENSRNOG00000021010 | ARL2 | 0.148157 | 0.037628 |
| ENSRNOG00000019009 | Arrdc2 | 0.312296 | 0.002981 |
| ENSRNOG00000014490 | BDH2 | -0.49399 | 0.039887 |
| ENSRNOG00000014448 | BMAL1 | -0.32112 | 0.009952 |
| ENSRNOG00000053384 | BMP7 | -0.97209 | 0.039943 |
| ENSRNOG00000015835 | CACNA2D2 | 0.24664 | 0.048015 |
| ENSRNOG00000025895 | CAVIN2 | -0.27997 | 0.026906 |
| ENSRNOG00000019586 | CCDC24 | 0.762861 | 0.049463 |
| ENSRNOG00000008697 | CCN3 | -0.62043 | 0.046887 |
| ENSRNOG00000025332 | CD109 | -0.6332 | 0.035651 |
| ENSRNOG00000010253 | CD163 | -0.72643 | 0.004776 |
| ENSRNOG00000008517 | CDC42EP1 | 0.208077 | 0.022031 |
| ENSRNOG00000020904 | CDC42EP2 | 0.230764 | 0.015401 |
| ENSRNOG00000040266 | CDKL4 | -0.46685 | 0.031964 |
| ENSRNOG00000053230 | CEP20 | -0.27037 | 0.003119 |
| ENSRNOG00000025012 | CHAT | 1.757314 | 0.041031 |
| ENSRNOG00000001915 | CHODL | -0.57933 | 0.003289 |
| ENSRNOG00000000572 | CHST3 | 0.387017 | 0.012454 |
| ENSRNOG00000019174 | CHTF18 | 0.466011 | 0.040427 |
| ENSRNOG00000012752 | CKLF | -0.86712 | 0.034926 |
| ENSRNOG00000012565 | CLN8 | 0.228103 | 0.015323 |
| ENSRNOG00000016828 | CMTM5 | 0.141117 | 0.039933 |
| ENSRNOG00000043151 | CNTLN | -0.29028 | 0.034474 |
| ENSRNOG00000027016 | COBLL1 | 0.245932 | 0.044513 |
| ENSRNOG00000005286 | Coch | -0.73687 | 0.037771 |
| ENSRNOG00000060008 | COG7 | 0.190667 | 0.033451 |
| ENSRNOG00000001249 | COL6A1 | -0.71385 | 0.02934 |
| ENSRNOG00000010841 | COL8A2 | -1.164 | 0.011937 |
| ENSRNOG00000016366 | COLEC12 | -0.72708 | 0.043306 |
| ENSRNOG00000017012 | COQ7 | 0.271053 | 0.026476 |
| ENSRNOG00000050052 | COX19 | 0.187165 | 0.029244 |
| ENSRNOG00000021220 | CPXM1 | 0.430207 | 0.031913 |
| ENSRNOG00000011207 | CSHL1 | -7.68787 | 8.96E-13 |
| ENSRNOG00000021155 | CTSK | -0.67043 | 0.027786 |
| ENSRNOG00000014479 | CTTNBP2NL | 0.197687 | 0.027058 |
| ENSRNOG00000015980 | DCLRE1C | 0.42685 | 0.004659 |
| ENSRNOG00000056106 | DIP2B | 0.189139 | 0.017578 |
| ENSRNOG00000055934 | Dmkn | 1.195792 | 0.034064 |
| ENSRNOG00000002947 | Dpt | -1.39837 | 0.017427 |
| ENSRNOG00000047905 | DUS1L | -0.1876 | 0.038314 |
| ENSRNOG00000006628 | DUSP16 | 0.234622 | 0.034412 |
| ENSRNOG00000010789 | DUSP7 | 0.210164 | 0.006753 |
| ENSRNOG00000018207 | DYNLT1 | 0.415377 | 0.019068 |
| ENSRNOG00000004449 | E2F6 | 0.192012 | 0.020394 |
| ENSRNOG00000009224 | E4F1 | 0.192518 | 0.048108 |
| ENSRNOG00000015596 | EFHD1 | 0.170401 | 0.036776 |
| ENSRNOG00000033979 | EIF1 | 0.471246 | 0.031683 |
| ENSRNOG00000031421 | EIF1AX | -0.17023 | 0.04371 |
| ENSRNOG00000017654 | EMC8 | -0.21293 | 0.016165 |
| ENSRNOG00000024364 | ENKD1 | 0.508195 | 0.033744 |
| ENSRNOG00000037340 | EPHA10 | 0.267151 | 0.015344 |
| ENSRNOG00000004964 | ERBB3 | 0.275329 | 0.004127 |
| ENSRNOG00000009343 | EVPL | -1.26114 | 0.008216 |
| ENSRNOG00000015957 | F13A1 | -0.82538 | 0.008448 |
| ENSRNOG00000007608 | FEZF1 | -0.79582 | 0.00348 |
| ENSRNOG00000012278 | FGF10 | -0.45737 | 0.027443 |
| ENSRNOG00000042753 | FGF13 | -0.24198 | 0.033442 |
| ENSRNOG00000007764 | FRMD4B | 0.188016 | 0.036491 |
| ENSRNOG00000016189 | FRMD5 | 0.200608 | 0.042934 |
| ENSRNOG00000016351 | FRRS1 | -0.54921 | 0.038849 |
| ENSRNOG00000018040 | GINS4 | -0.39577 | 0.049411 |
| ENSRNOG00000038328 | GJC2 | 0.254145 | 0.038624 |
| ENSRNOG00000000138 | GLIS1 | -0.77584 | 0.039369 |
| ENSRNOG00000001235 | GNA12 | 0.124575 | 0.035582 |
| ENSRNOG00000011335 | GPR50 | -0.76728 | 0.026798 |
| ENSRNOG00000005519 | GRM3 | 0.262863 | 0.044967 |
| ENSRNOG00000042905 | H2-T24 | 0.297421 | 0.020584 |
| ENSRNOG00000018870 | HAPLN2 | 0.200186 | 0.035415 |
| ENSRNOG00000026661 | HCAR1 | -1.81262 | 0.003118 |
| ENSRNOG00000037799 | HDX | 0.60565 | 0.040871 |
| ENSRNOG00000023969 | HERC6 | 0.379095 | 0.004456 |
| ENSRNOG00000001338 | HPD | 0.852897 | 0.020555 |
| ENSRNOG00000011427 | HR | 0.195942 | 0.012993 |
| ENSRNOG00000049994 | IFI44L | 0.625514 | 0.03903 |
| ENSRNOG00000006227 | IFIH1 | 0.386278 | 0.020164 |
| ENSRNOG00000022839 | IFIT3 | 0.797305 | 0.004834 |
| ENSRNOG00000021966 | IL17RD | 0.433556 | 0.020998 |
| ENSRNOG00000014083 | IQSEC3 | 0.248028 | 0.028772 |
| ENSRNOG00000017414 | IRF7 | 0.465709 | 0.043758 |
| ENSRNOG00000004516 | ITGBL1 | -0.63393 | 0.043903 |
| ENSRNOG00000027590 | JAKMIP3 | 0.152313 | 0.037616 |
| ENSRNOG00000016467 | KCTD1 | -0.23868 | 0.036702 |
| ENSRNOG00000031269 | KIAA0930 | 0.151568 | 0.024091 |
| ENSRNOG00000018168 | KLC4 | 0.212783 | 0.019482 |
| ENSRNOG00000018867 | KLHDC7A | -0.4054 | 0.030854 |
| ENSRNOG00000003289 | LAP3 | 0.134421 | 0.049479 |
| ENSRNOG00000006865 | LAPTM4A | -0.16247 | 0.039383 |
| ENSRNOG00000018427 | LHX3 | -1.28223 | 0.024998 |
| ENSRNOG00000028348 | LHX8 | 3.178332 | 0.043962 |
| ENSRNOG00000055952 | LOC120096422 | -0.6086 | 0.035456 |
| ENSRNOG00000058132 | LOC120100227 | 0.425724 | 0.048938 |
| ENSRNOG00000012952 | LRIG1 | 0.142252 | 0.02727 |
| ENSRNOG00000048725 | LSM2 | 0.325648 | 0.047063 |
| ENSRNOG00000010642 | LYSMD2 | 0.176128 | 0.032214 |
| ENSRNOG00000021023 | MAG | 0.19972 | 0.028224 |
| ENSRNOG00000009113 | MARCKSL1 | 0.296248 | 0.026371 |
| ENSRNOG00000016516 | Mbp | 0.173999 | 0.039233 |
| ENSRNOG00000013790 | MFSD1 | -0.14758 | 0.039779 |
| ENSRNOG00000016038 | MGMT | 0.710516 | 0.014119 |
| ENSRNOG00000046171 | MLXIPL | 0.404174 | 0.007868 |
| ENSRNOG00000000775 | MOG | 0.16753 | 0.03768 |
| ENSRNOG00000014060 | MRAS | -0.16871 | 0.011612 |
| ENSRNOG00000020991 | MS4A6A | -0.85835 | 0.019382 |
| ENSRNOG00000001963 | MX1 | 0.323251 | 0.026657 |
| ENSRNOG00000055858 | Myb | 0.503305 | 0.028747 |
| ENSRNOG00000025757 | MYH6 | 0.208649 | 0.021739 |
| ENSRNOG00000016256 | MYO9B | 0.159496 | 0.026749 |
| ENSRNOG00000028274 | MYRF | 0.206054 | 0.027367 |
| ENSRNOG00000053875 | Nacad | 0.157724 | 0.016068 |
| ENSRNOG00000054157 | NADK2 | -0.13616 | 0.025394 |
| ENSRNOG00000019031 | NEU4 | 0.693142 | 0.045367 |
| ENSRNOG00000005392 | NGFR | 0.796118 | 0.016712 |
| ENSRNOG00000010282 | NINJ2 | 0.328295 | 0.048133 |
| ENSRNOG00000006255 | NIPAL4 | 0.33034 | 0.018282 |
| ENSRNOG00000003794 | NMRAL1 | 0.277265 | 0.043106 |
| ENSRNOG00000003741 | NPTX1 | -0.29692 | 0.000465 |
| ENSRNOG00000001006 | NPTX2 | -0.48095 | 0.047836 |
| ENSRNOG00000003765 | NR0B1 | -1.30236 | 0.039233 |
| ENSRNOG00000017420 | NUDT6 | -0.31024 | 0.03496 |
| ENSRNOG00000017919 | NUP133 | 0.257156 | 0.04646 |
| ENSRNOG00000001369 | OAS1 | 0.629395 | 0.026379 |
| ENSRNOG00000028814 | Oasl2 | 0.350816 | 0.042683 |
| ENSRNOG00000013263 | ODAD3 | 0.685444 | 0.018562 |
| ENSRNOG00000014034 | OLFML2A | -0.34168 | 0.03321 |
| ENSRNOG00000028648 | OLIG1 | 0.169699 | 0.014267 |
| ENSRNOG00000028658 | OLIG2 | 0.208414 | 0.024477 |
| ENSRNOG00000047517 | Oxld1 | -0.38478 | 0.01518 |
| ENSRNOG00000033479 | PCDHB11 | 0.666737 | 0.028031 |
| ENSRNOG00000019005 | PDE8A | 0.216307 | 0.032739 |
| ENSRNOG00000026985 | PHLDB1 | 0.174265 | 0.012786 |
| ENSRNOG00000000274 | PHYHIPL | 0.145613 | 0.04086 |
| ENSRNOG00000039850 | PIGP | -0.29923 | 0.007132 |
| ENSRNOG00000000529 | PIM1 | -0.51857 | 0.043639 |
| ENSRNOG00000029698 | PIM3 | 0.20328 | 0.009014 |
| ENSRNOG00000006570 | PLEKHG3 | 0.155927 | 0.026533 |
| ENSRNOG00000010650 | PLEKHH1 | 0.242468 | 0.033022 |
| ENSRNOG00000013751 | PLPBP | -0.15558 | 0.026106 |
| ENSRNOG00000020906 | POLA2 | 0.38312 | 0.025333 |
| ENSRNOG00000047686 | POU3F1 | 0.671601 | 0.024854 |
| ENSRNOG00000004719 | PP2D1 | -0.29884 | 0.024322 |
| ENSRNOG00000027408 | PPID | -0.13611 | 0.042954 |
| ENSRNOG00000009882 | PPP3CA | -0.18402 | 0.046992 |
| ENSRNOG00000030486 | PRDM6 | -2.46488 | 0.003731 |
| ENSRNOG00000008915 | PRIMA1 | 0.344246 | 0.042394 |
| ENSRNOG00000017374 | PRL | -4.48017 | 7.37E-13 |
| ENSRNOG00000004160 | PRPS2 | -0.26239 | 0.021708 |
| ENSRNOG00000060193 | PRR29 | 0.640691 | 0.010223 |
| ENSRNOG00000004666 | PRR5L | 0.313416 | 0.005572 |
| ENSRNOG00000057616 | Ptch2 | 0.332081 | 0.019785 |
| ENSRNOG00000003253 | QDPR | 0.166368 | 0.047732 |
| ENSRNOG00000028872 | RAI14 | -0.39225 | 0.018151 |
| ENSRNOG00000010219 | RALGDS | 0.157176 | 0.021637 |
| ENSRNOG00000052173 | RANBP3L | -0.72162 | 0.042335 |
| ENSRNOG00000059961 | RAPGEF3 | 0.173005 | 0.010205 |
| ENSRNOG00000002097 | RASL11B | 0.403947 | 0.046015 |
| ENSRNOG00000026295 | RBPJL | 0.370355 | 0.004586 |
| ENSRNOG00000016309 | RGP1 | 0.249586 | 0.024283 |
| ENSRNOG00000005911 | RLN3 | 2.516892 | 0.003812 |
| ENSRNOG00000027017 | RNASEL | -0.19969 | 0.04734 |
| ENSRNOG00000008807 | RP1 | 1.004403 | 0.02393 |
| ENSRNOG00000018812 | RPP25 | 0.38003 | 0.019059 |
| ENSRNOG00000054493 | Rpph1 | 1.10439 | 0.038926 |
| ENSRNOG00000017309 | RSRP1 | 0.130232 | 0.006689 |
| ENSRNOG00000000636 | RTKN2 | 0.360433 | 0.049238 |
| ENSRNOG00000004672 | SEC14L2 | 0.160263 | 0.024884 |
| ENSRNOG00000002723 | SELE | 1.24884 | 0.012475 |
| ENSRNOG00000016254 | SEMA4C | 0.167745 | 0.041663 |
| ENSRNOG00000013679 | SEMA4D | 0.162862 | 0.02034 |
| ENSRNOG00000011763 | SERP1 | -0.14772 | 0.04705 |
| ENSRNOG00000018412 | SFI1 | 0.286572 | 0.014163 |
| ENSRNOG00000020726 | SIPA1 | 0.268334 | 0.006923 |
| ENSRNOG00000021644 | SLC15A3 | 0.533654 | 0.033821 |
| ENSRNOG00000018131 | SLC16A4 | -0.47151 | 0.023886 |
| ENSRNOG00000029465 | Slc26a10 | 0.532222 | 0.006231 |
| ENSRNOG00000006096 | SLC26A7 | -1.40454 | 0.046559 |
| ENSRNOG00000011723 | SLC44A3 | -0.96203 | 0.009299 |
| ENSRNOG00000009005 | SLCO2A1 | -0.86988 | 0.041944 |
| ENSRNOG00000032798 | SLCO3A1 | 0.185224 | 0.049084 |
| ENSRNOG00000009209 | SLITRK1 | -0.1577 | 0.04178 |
| ENSRNOG00000018359 | SMAD7 | 0.188173 | 0.028613 |
| ENSRNOG00000019536 | SMIM3 | -0.34019 | 0.042557 |
| ENSRNOG00000001056 | SNAPC2 | 0.221476 | 0.00342 |
| ENSRNOG00000061138 | Snord30 | -0.77413 | 0.033696 |
| ENSRNOG00000018841 | SOX8 | 0.227099 | 0.01772 |
| ENSRNOG00000025787 | SPAG6 | 0.474152 | 0.022078 |
| ENSRNOG00000003463 | SREBF1 | 0.155593 | 0.017985 |
| ENSRNOG00000007998 | SSB | -0.14673 | 0.024793 |
| ENSRNOG00000011148 | SSR3 | -0.13703 | 0.048697 |
| ENSRNOG00000019294 | STK16 | 0.133323 | 0.034738 |
| ENSRNOG00000033389 | SUSD2 | -0.16108 | 0.04874 |
| ENSRNOG00000030625 | TF | 0.214073 | 0.023335 |
| ENSRNOG00000045829 | THBS1 | -0.70777 | 0.005293 |
| ENSRNOG00000004303 | TIMP3 | -0.22527 | 0.028476 |
| ENSRNOG00000012860 | TMEM184C | -0.15602 | 0.036281 |
| ENSRNOG00000048231 | TMEM235 | 0.688469 | 0.008661 |
| ENSRNOG00000026136 | TNFAIP8 | -0.26248 | 0.041557 |
| ENSRNOG00000028041 | TNNT1 | 0.763989 | 0.014623 |
| ENSRNOG00000033734 | TNNT2 | -1.63169 | 0.035975 |
| ENSRNOG00000050500 | TOB2 | 0.279199 | 0.00527 |
| ENSRNOG00000029141 | TRABD2B | -1.33029 | 0.013641 |
| ENSRNOG00000004110 | Trib2 | -0.17556 | 0.042425 |
| ENSRNOG00000002469 | TRIM7 | 0.53498 | 0.024941 |
| ENSRNOG00000058681 | TSGA10 | 0.201338 | 0.046027 |
| ENSRNOG00000056362 | Ttll13 | 0.439293 | 0.002677 |
| ENSRNOG00000058586 | TYRO3 | 0.167836 | 0.018763 |
| ENSRNOG00000009345 | UGT8 | 0.227198 | 0.013751 |
| ENSRNOG00000000567 | UNC5B | 0.252441 | 0.009036 |
| ENSRNOG00000029071 | UNC5C | -0.16463 | 0.028205 |
| ENSRNOG00000028456 | USH1G | 0.604898 | 0.046594 |
| ENSRNOG00000027012 | USP54 | 0.170988 | 0.024834 |
| ENSRNOG00000005823 | UTP20 | 0.194945 | 0.034388 |
| ENSRNOG00000047457 | VIPR1 | -1.15253 | 0.024172 |
| ENSRNOG00000008668 | VSIG8 | 0.986069 | 0.038215 |
| ENSRNOG00000028783 | WDR35 | -0.17339 | 0.005655 |
| ENSRNOG00000013336 | WDR70 | -0.31609 | 0.002238 |
| ENSRNOG00000025735 | Wdr86 | -1.47561 | 0.026507 |
| ENSRNOG00000007869 | WSCD1 | 0.174259 | 0.02959 |
| ENSRNOG00000014564 | YIPF5 | -0.16957 | 0.020063 |
| ENSRNOG00000056617 | ZSWIM8 | 0.555607 | 0.033258 |
| ENSRNOG00000025818 | ZXDC | 0.216438 | 0.038315 |

**Table S7.** Significantly differentially expressed genes (DEGs) between Study 1 and Study 2 *Krtcap3* knock-out (KO) rats.
